# Persistent acetylation of histone H3 lysine 56 compromises the activity of DNA replication origins

**DOI:** 10.1101/2022.03.22.485257

**Authors:** Roch Tremblay, Yosra Mehrjoo, Antoine Simoneau, Mary E. McQuaid, Corey Nislow, Guri Giaever, Hugo Wurtele

**Affiliations:** Maisonneuve-Rosemont Hospital Research Center, 5415 blvd L’assomption, Montreal, H1T 2M4, Canada; Molecular Biology Program, Université de Montréal, 2900 Edouard-Montpetit, Montreal, Québec, Canada, H3T 1J4; Department of Pharmaceutical Sciences, University of British Columbia, Vancouver, Canada; Department of Medicine, Université de Montréal, 2900 Edouard-Montpetit, Montreal, Québec, Canada, H3T 1J4

## Abstract

In *Saccharomyces cerevisiae*, newly synthesized histone H3 are acetylated on lysine 56 (H3 K56ac) by the Rtt109 acetyltransferase prior to their deposition on nascent DNA behind replication forks. Two deacetylases of the sirtuin family, Hst3 and Hst4, remove H3 K56ac from chromatin following S phase. *hst3*Δ *hst4*Δ cells present constitutive H3 K56ac, which sensitizes cells to replicative stress via mechanisms that remain unclear. We performed a screen to identify genes that influence cell fitness upon nicotinamide (NAM)-induced inhibition of sirtuins. The screen revealed that *DBF4* heterozygosity causes NAM sensitivity. *DBF4* and *CDC7* encode subunits of the Dbf4-dependent kinase, which activates origins of DNA replication. We show that i) cells harboring the *dbf4-1* or *cdc7-4* hypomorphic alleles are sensitive to NAM, ii) Rif1, an inhibitor of Cdc7-dependent activation of origins, causes DNA damage and replication defects in NAM-treated cells and *hst3*Δ *hst4*Δ mutants, and iii) *cdc7-4 hst3*Δ *hst4*Δ cells display synthetic temperature sensitivity associated with delayed initiation of DNA replication. Such replication defects are not due to activation of the intra-S phase checkpoint but require Rtt109-dependent H3 K56ac. Overall, these results suggest that persistent H3 K56ac sensitizes cells to replicative stress in part by negatively influencing replication origin activity.

## INTRODUCTION

DNA replication initiates at multiple origins throughout chromosomes during the S phase of the cell cycle (1). During G1, Cdt1 and Cdc6 load the MCM helicase complex on DNA at origins of replication bound by the Origin Recognition Complex (ORC). At the beginning of S phase, cyclin-dependent (CDK) and Dbf4-dependent (DDK) kinase activities promote the recruitment of factors including Cdc45 and the GINS complex to replication origins as well as the activation of the MCM helicase. Melting of origin DNA resulting from MCM helicase activity allows the formation of two replication forks (RF) that travel in opposite directions along chromosomal DNA. Depending on the timing of their activation in S phase, eukaryotic origins are classified as early, mid, or late. Such sequential activation of origins has been shown to result at least in part from the recycling of limiting replication initiation factors from early to mid, and then to late replicating genomic regions (2, 3). Such temporal organization of DNA replication is evolutionarily conserved among eukaryotes; however, the repertoire of cellular factors and molecular mechanisms modulating origin activation remains incompletely characterized.

RF progression can be halted upon encountering DNA lesions induced by any among a multitude of environmental or endogenous genotoxins (4). This engenders a state of replicative stress which can prevent completion of chromosomal duplication, thereby causing genomic instability. Stalled RF activate Mec1 (ATR in humans), the apical kinase of the intra-S phase checkpoint response in yeast (4). In turn, Mec1 promotes activation of the kinase Rad53 via one of two pathways that depend upon either the RF component Mrc1 or the adaptor protein Rad9 (5). Activated Mec1 and Rad53 phosphorylate a plethora of substrates to i) promote the stability of stalled RF, and ii) to prevent further activation of replication origins (6). In yeast, this latter effect has been shown to depend on Rad53-dependent phosphorylation of the key replication factors Dbf4 and Sld3, which prevents activation of MCM helicase complexes at origins that have not yet been fired (7). Intra-S phase checkpoint-dependent inhibition of origin activity is important to prevent inordinate accumulation and eventual collapse of stalled RF during periods of genotoxic stress (8).

Histone post-translational modifications are critical determinants of DNA replication dynamics and origin activity (9, 10). Among those modifications, histone lysine acetylation can either promote and inhibit origin activity depending on the identity of the modified residue and/or chromosomal context (11). The sirtuin family of histone deacetylases is well-conserved throughout evolution, and several of its members have been shown to influence DNA replication and repair (12). The yeast *Saccharomyces cerevisiae* possesses 5 sirtuins: the founding member, Sir2, and Homologues of Sir Two 1 through 4 (Hst1-4) (12). Sir2 targets histone H4 lysine 16 acetylation (H4 K16ac), which regulates origins at the rDNA locus and telomeres (13, 14). Hst1, which can also target H4 K16ac, forms a complex with Sum1 and Rfm1 and modulates the activity of a subset of origins genome-wide (15, 16). While the impact of Hst2 on DNA replication has not been directly assessed, at least some of the functions of this sirtuin are known to be partially redundant with those of Sir2, as overexpressed Hst2 rescues gene silencing defects caused by *sir2*Δ (17, 18).

The only known histone substrate of the redundant sirtuins Hst3 and Hst4 is acetylated H3 lysine 56 (H3 K56ac) (19). This modification is catalyzed by the acetyltransferase Rtt109 on newly synthesized histones H3 prior to their deposition onto nascent DNA during S phase (20, 21). After S phase, Hst3 and Hst4 remove H3 K56ac chromosome-wide such that a large majority of nucleosomes do not harbor H3 K56ac at the start of the next cell cycle. While under normal circumstances the bulk of H3 K56ac is removed by Hst3 during G2, Hst4 can compensate for its absence. As such, the stoichiometry of H3 K56ac approaches 100% throughout the cell cycle in *hst3*Δ *hst4*Δ double mutants (19, 22). While constitutive H3 K56ac has been shown to cause spontaneous DNA damage, thermosensitivity, and increased sensitivity to genotoxins that cause replicative stress (19, 23, 24), the molecular mechanisms underlying such striking phenotypes remain poorly understood.

Nicotinamide (NAM) is a non-competitive pan-inhibitor of sirtuins (25). Our group previously performed genetic screens in *S. cerevisiae* with the goal of identifying genes whose homozygous deletion (i.e. complete loss-of-function) confers either fitness defect or advantage in response to NAM-induced sirtuin inhibition and consequent H3 K56 hyperacetylation (26). These screens revealed that several genes encoding regulators of the DNA replication stress response promote resistance to NAM-induced elevation in H3 K56ac caused by inhibition of Hst3 and Hst4 (26, 27). Previously published data also indicate that cells lacking HST3 are defective in the maintenance of artificial chromosomes harboring a reduced number of DNA replication origins (28), further linking H3 K56ac with the regulation of DNA replication dynamics. Here, we present the results of a genome-wide screen aimed at identifying genes whose haploinsufficiency modulates cell fitness in response to NAM. Overall, we found that i) appropriate dosage of genes involved in various cellular pathways influence cell fitness in response to NAM, ii) factors promoting DNA replication origin activation are critical for survival in the absence of Hst3 and Hst4 activity, and ii) abnormal persistence of the acetylation of new histones H3 on lysine 56 throughout the cell cycle compromises the activity of replication origins.

## RESULTS

### A genetic screen to identify genes modulating cellular fitness in response to NAM

We performed a screen using the pooled yeast strains of the heterozygote diploid collection to identify haploinsufficient genes that influence cell fitness upon NAM exposure (Table S1). Using a Z-score cut-off of +/- 2.58 (99% cumulative percentage), the screen identified 131 and 58 genes whose heterozygosity caused reduced or increased fitness, respectively, during propagation for 20 generations in YPD medium containing 41 mM NAM (Figure 1A). This list of genes presents only modest overlap with that obtained from our previously published screen using the homozygote deletion strain collection (Figure 1B), suggesting that most of the genes identified in the latter screen are not haploinsufficient with regards to NAM sensitivity. We note that such limited overlap between screens performed on the homozygous and heterozygous deletion collections has also been observed in several other chemogenetic screens (29). Gene Ontology (GO) term analysis of genes whose heterozygosity sensitizes cells to NAM revealed an obvious enrichment in DNA replication and DNA damage response pathway, whereas terms reflecting proteasome-related and catabolic processes were associated with mutations that enhanced fitness in NAM (Figure 1C and Table S2-S3).

**FIGURE 1.**
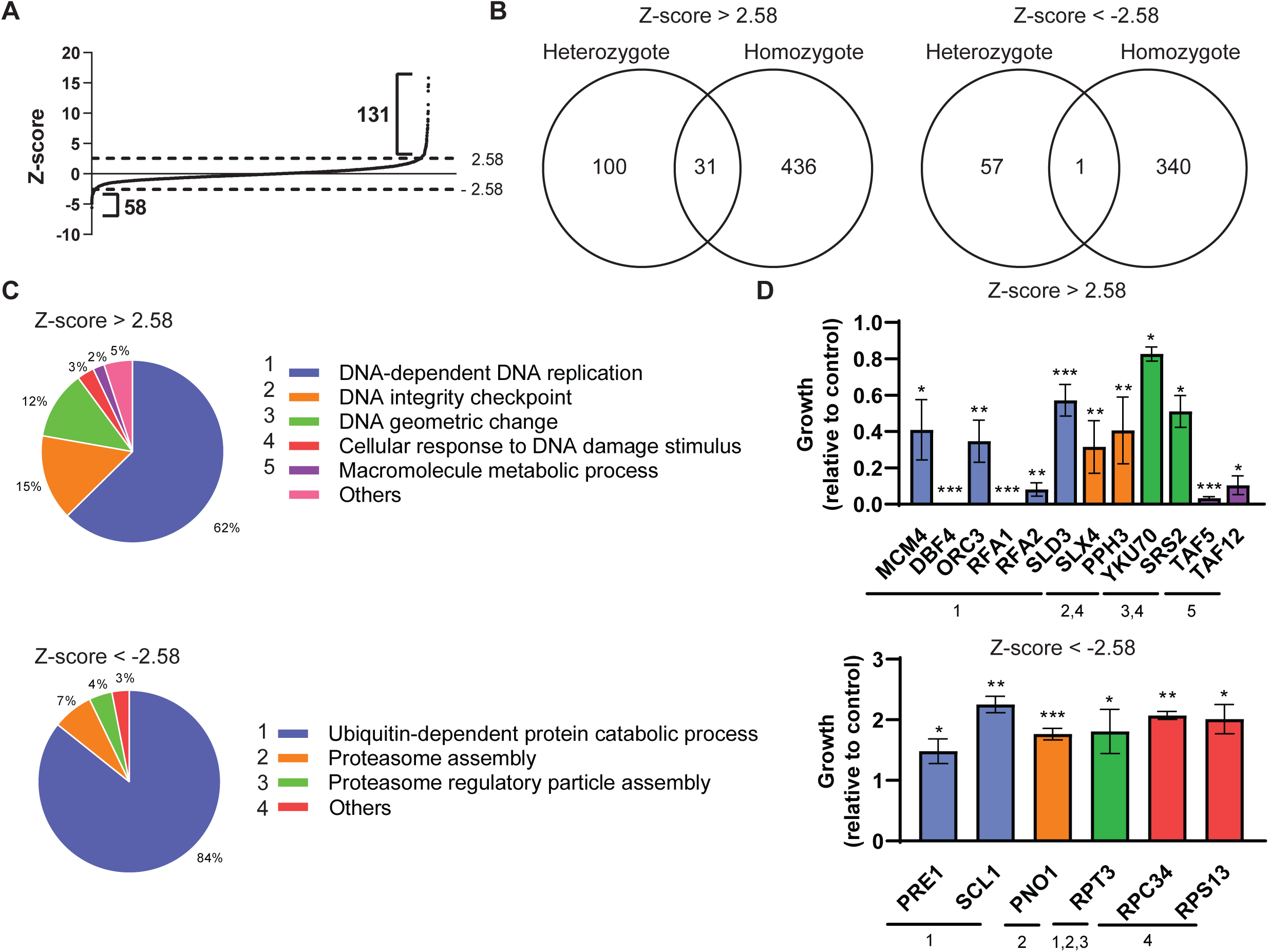
A chemogenomic screen identifies genes whose heterozygosity modulates cell fitness upon NAM exposure. (A) Z-score of individual heterozygote diploid yeast strains after 20 generations in medium containing 41mM NAM. GO-terms of genes for which the Z-scores is > 2.58 or < -2.58 were further analyzed in B and C. (B) Venn diagram comparing the heterozygote diploid screen presented in this article with a previous one performed with homozygote diploid deletion strains (26). (C) GO-term associated with genes presenting Z-scores > 2.58 or < -2.58. (D) Growth competition assays for selected heterozygote deletion strains presenting Z-scores > 2.58 or < -2.58. WT and mutant cells were mixed 1:1 and grown in YPD +/- 41 mM NAM for 20 generations. The fraction of mutant/WT cells in the culture was assessed by plating on selective medium followed by colony counting. Colors and numbers below strain names refer to GO-terms in C.

We next sought to validate individual heterozygous mutations representing the main categories of “hits’’ identified in the screen. WT diploid and heterozygote mutant strains of interest (*yfg1*Δ::KanMX*/YFG1)* were mixed in a 1:1 ratio and incubated for 20 generations in YPD +/- NAM. Appropriate dilutions of cells were then plated on YPD-agar +/- G418, and the ratio of the number of heterozygous mutant (G418-resistant) vs WT (G418-sensitive) colonies was calculated (Figure 1D). These competition assays confirmed the expected impact of heterozygous mutations causing diminished cell fitness in NAM-containing medium, thereby validating our screen results. While significant improvement in growth was observed for individual heterozygous mutants expected to promote fitness in NAM, we note that heterozygous mutations causing improved fitness in response to NAM displayed generally lower absolute Z-scores than those reducing fitness (Figure 1A, Table S1).

### Reduced activity DNA replication origins activity sensitizes cells to NAM

As mentioned previously, cells lacking Hst3 have previously been shown to present defects in the maintenance of an artificial chromosome harboring reduced number of DNA replication origins, revealing a potential link between this sirtuin and origin activity (28). Nevertheless, the mechanistic basis explaining the effect of Hst3 on origins remains unknown. Interestingly, 6 of the 11 essential DNA replication genes identified in the screen as promoting NAM resistance are members of the pre-replicative complex (ORC3 and MCM4) or involved in various steps of origin activation (DBF4, SLD2, SLD3, PSF2). We therefore decided to further investigate the possible relationship between NAM sensitivity and origin activity. To this end, we focused on *DBF4* and its associated kinase Dbf4-dependent kinase (DDK), a complex formed by Dbf4 and the Cdc7 kinase. Even though *CDC7* was not identified as being haploinsufficient with regards to NAM resistance in our screen, haploid cells expressing hypomorphic temperature sensitive alleles of *DBF4* (*dbf4-1*) and *CDC7* (*cdc7-4*) were found to be NAM-sensitive at 30°C, a semi-permissive temperature for these alleles (Figure 2A). Our data also indicate that *bob1-1 cdc7*Δ cells, which harbor a mutation in *MCM5* that bypasses DDK-dependent phosphorylation of the MCM complex that is necessary for origin activation (30), are not sensitive to NAM (Figure 2B). This argues that the MCM complex is likely to be the relevant target of DDK in this context, and suggests that impaired activation of replication origins sensitizes cells to NAM.

**FIGURE 2.**
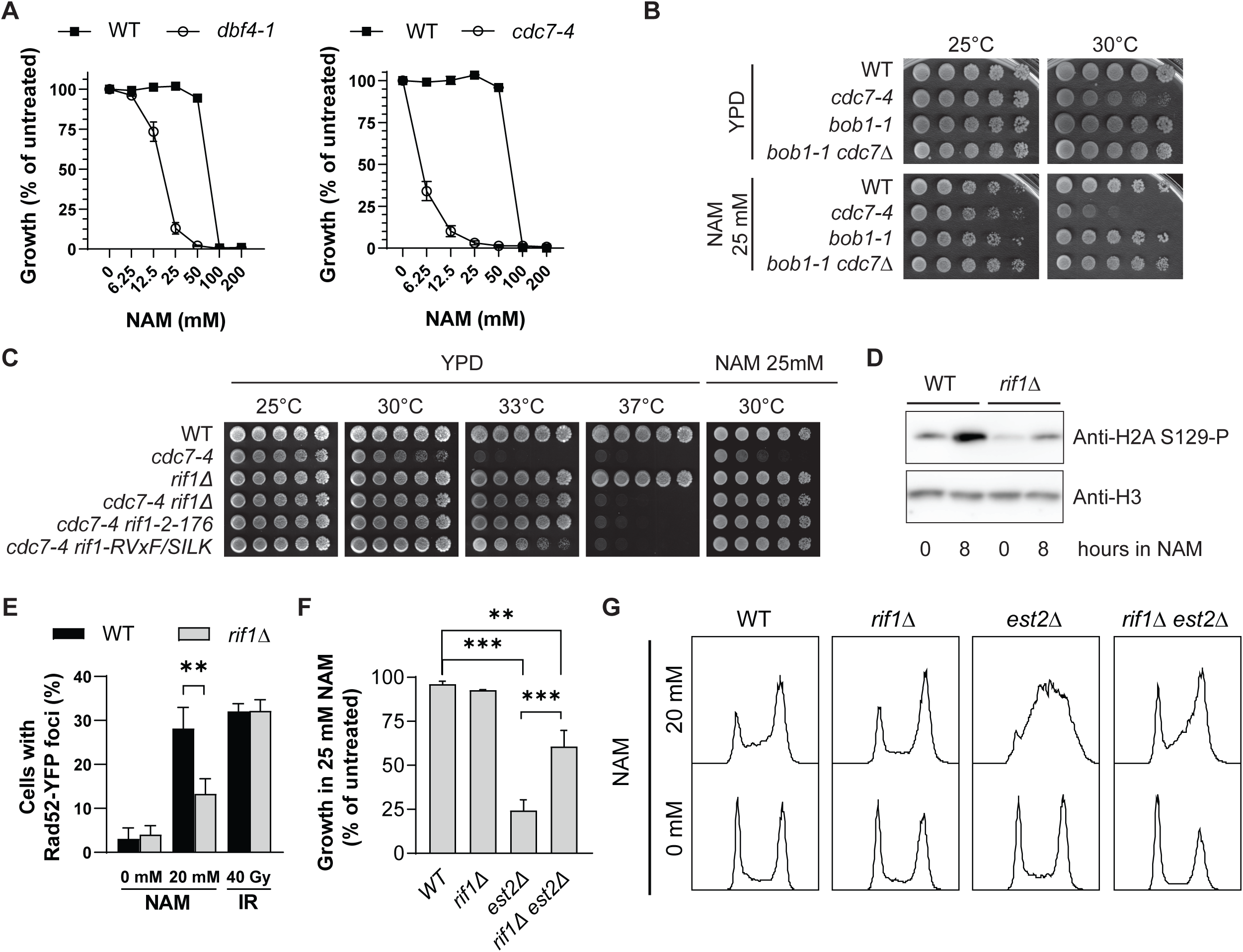
Yeast cells harboring hypomorphic alleles of *CDC7* or *DBF4* are sensitive to NAM. (A) Haploid WT, *dbf4-1* and *cdc7-4* cells were incubated at 30°C in medium containing the indicated concentration of NAM. OD_630_ readings were taken at 72 h to evaluate cell proliferation (see Methods). (B) 5-fold serial dilutions of cell cultures were spotted on YPD-agar and YPD-agar + 25 mM NAM plates. Plates were incubated at the indicated temperature. (C) As in B. (D) WT and *rif1*Δ cells were exposed to 20 mM NAM for 8 h at 30°C before harvest for immunoblotting. (E) Exponentially growing WT and *rif1*Δ cells in SC medium were treated with 20 mM NAM for 8 h at 30°C, or exposed to 40 Gy of ionizing radiations followed by incubation for 1 h at 30°C. Samples were then taken for fluorescence microscopy. Graph bars represent mean ± SEM of three independent experiments. (F) The ratio of OD_630_ readings of cells treated with 25 mM NAM vs incubated in control medium after 48 h of growth in 96-wells plates is presented. Graph bars represent mean ± SEM of three independent experiments each containing 4 technical replicates. (G) Exponentially growing cells were incubated at 25°C in YPD containing 20 mM NAM for 8 h. Samples were taken for flow cytometry analysis of DNA content. **: p < 0.01 and ***: p < 0.001, unpaired two-tailed Student’s *t*-test.

Rap1-Interacting Factor 1 (Rif1) acts in concert with the phosphatase Glc7 to reverse DDK-dependent MCM phosphorylation, thereby inhibiting origin activation (31–34). Moreover, a previous screen performed by our group identified *RIF1* among the few yeast genes whose homozygous deletion improved cell fitness in NAM-containing medium (26). We found that NAM-induced growth defects and accumulation of cells in S phase is rescued by deletion of *RIF1* (Figure 2C, 2G). Moreover, N-terminal truncation or mutations in the Glc7-interacting motif (*rif1*-RVxF/SILK) of Rif1, both of which were previously shown to partially suppress the temperature sensitivity of *cdc7-4* mutants by eliminating Rif1 binding to Glc7 (33, 35), also suppressed the NAM sensitivity of *cdc7-4* cells (Figure 2C). Overall, these data indicate that Rif1/Glc7-dependent dephosphorylation of MCM influences NAM sensitivity.

We and others previously showed that NAM treatment causes replicative stress and DNA damage in yeast (19, 26, 27). Since lack of MCM phosphorylation by DDK causes sensitivity to replicative stress-inducing drugs (36, 37), we tested the impact of *RIF1* deletion on NAM-induced DNA damage. Compared to WT, *rif1*Δ cells presented reduced NAM-induced histone H2A S129 phosphorylation and Rad52-YFP foci formation (Figure 2D-E), two well-known markers of replicative stress-induced DNA damage (38, 39). Importantly, lack of Rif1 did not compromise the formation of ionizing radiation (IR)-induced Rad52 foci, which are not primarily caused by replication-associated DNA lesions. We note that, in addition to its role in regulating DNA replication, Rif1 is known to limit telomere length by inhibiting telomerase activity (40). Moreover we previously showed that cells with short telomeres are sensitive to NAM-induced sirtuin inhibition (27); we therefore considered the possibility that abnormal telomere elongation in *rif1*Δ cells might favor NAM resistance. Contrary to this notion, deletion of *RIF1* suppressed NAM-induced growth and S phase progression defects in telomerase-defective *est2*Δ cells (Figure 2F-G), indicating that the role of Rif1 in modulating NAM sensitivity is independent of its influence on telomerase activity.

### Rif1 and Cdc7 influence the phenotypes of *hst3*Δ *hst4*Δ cells

We previously showed that NAM-induced DNA damage is attributable in large part to inhibition of Hst3 and Hst4, leading to elevated H3 K56ac (26). We found that deletion of *RIF1* rescued the temperature sensitivity of *hst3*Δ *hst4*Δ cells as well as the synthetic lethality of *hst3*Δ *hst4*Δ *sir2*Δ without noticeably affecting H3 K56ac levels (Figure 3A-B). We note that for unknown reasons *hst3*Δ *hst4*Δ cells are temperature sensitive in S288C-derived genetic backgrounds but not in W303 (our unpublished observations; e.g., compare Figure 3A and 3G). Because of this, while most of the experiments involving *hst3*Δ *hst4*Δ were done in W303-derived strains, certain experiments including the one presented in Figure 3A were done in the BY4741 background (Table 1 indicates the yeast strains used in each figure of this study).

**FIGURE 3.**
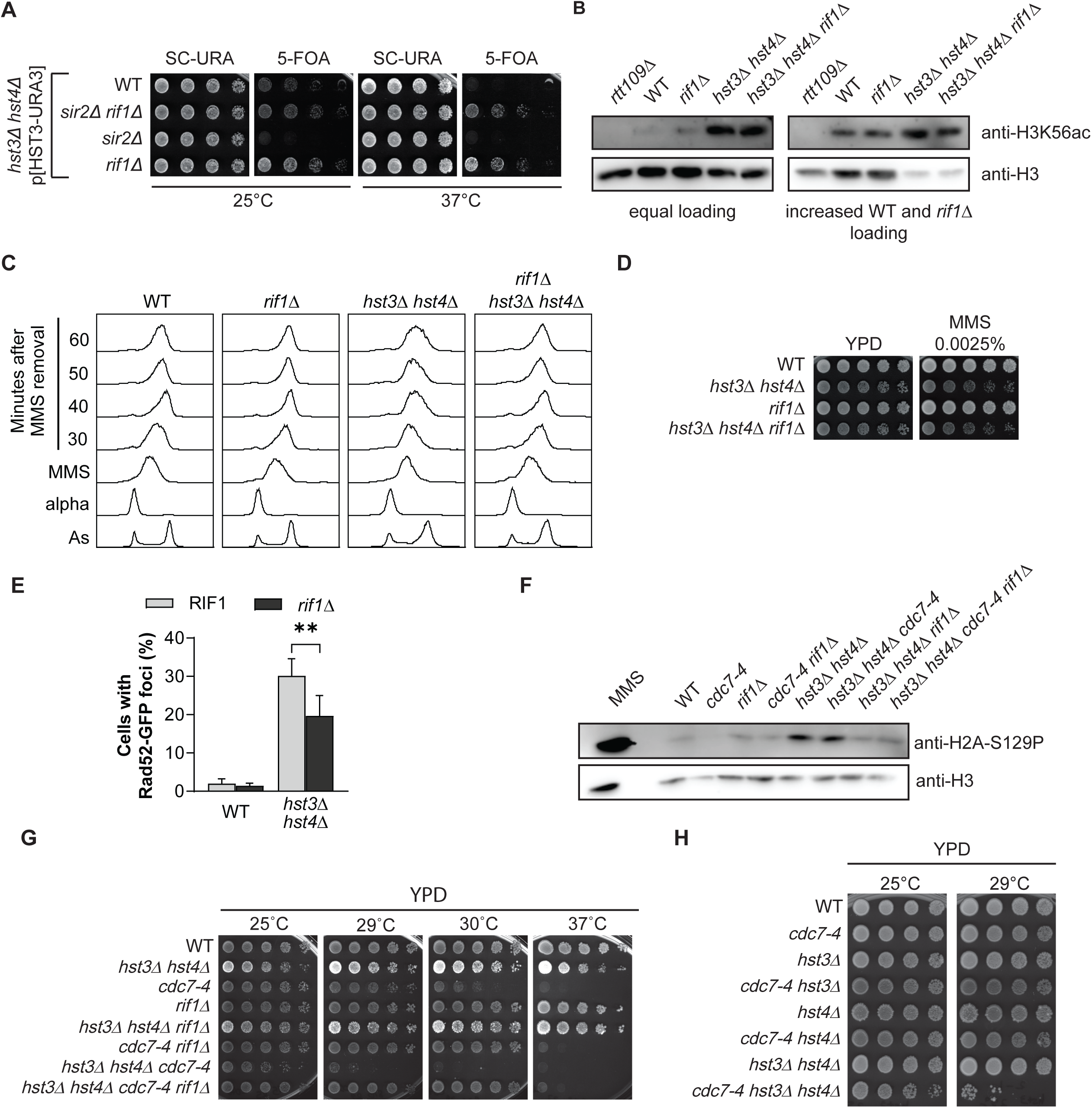
*rif1*Δ and *cdc7-4* influence the phenotypes of *hst3*Δ *hst4*Δ cells in opposite ways. (A) 5-fold serial dilutions of cell cultures were spotted on solid SC-URA and 5-FOA media. Plates were incubated at the indicated temperature. (B) Exponentially growing cells in YPD at 25°C were harvested and processed for immunoblotting. (C) Cells were arrested in G1 at 25°C using alpha factor (alpha) and released toward S in medium containing 0.01% of MMS for 90 minutes (MMS) at the same temperature. After MMS inactivation using sodium thiosulfate, cells were released in fresh YPD. Samples were taken for flow cytometry analysis of DNA content at the indicated time points. As: asynchronous. (D) 5-fold serial dilutions of cell cultures were spotted on YPD-agar and YPD-agar containing 0.0025 % MMS plates. (E) The fraction of cells harboring Rad52-GFP foci was assessed by fluorescence microscopy in exponentially growing cells at 25°C. Graph bars represent the mean value ± SEM of 10 independent cultures. **: p < 0.01, unpaired two-tailed Student’s *t-*test. (F) Exponentially growing cells in YPD at 25°C were harvested for immunoblotting. MMS: Cells were exposed to 0.03% MMS for 1 h prior to harvesting as control. (G) 5-fold serial dilutions of cell cultures were spotted on YPD-agar plates and incubated at the indicated temperature. (H) As in G.

**TABLE 1.**
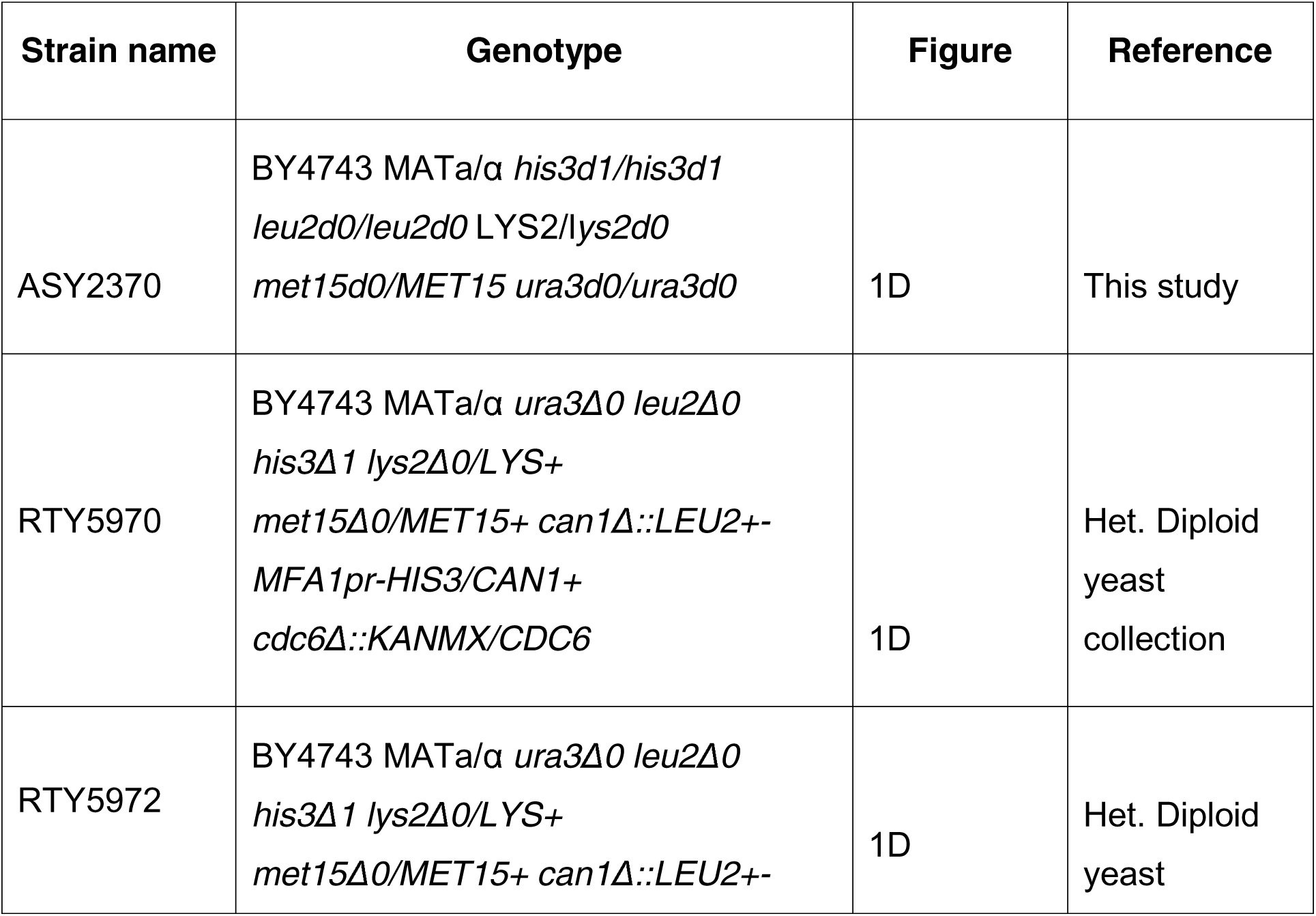
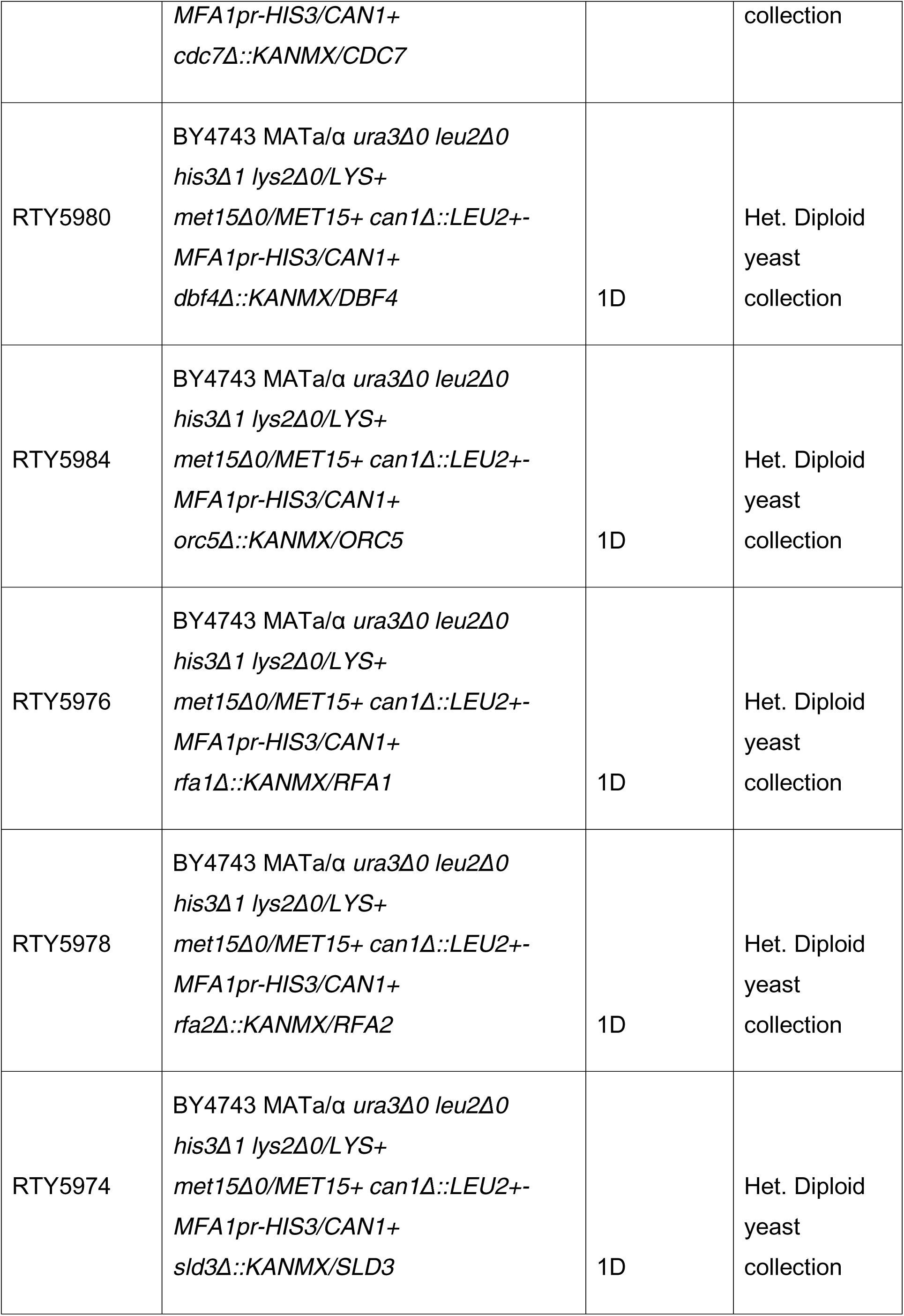
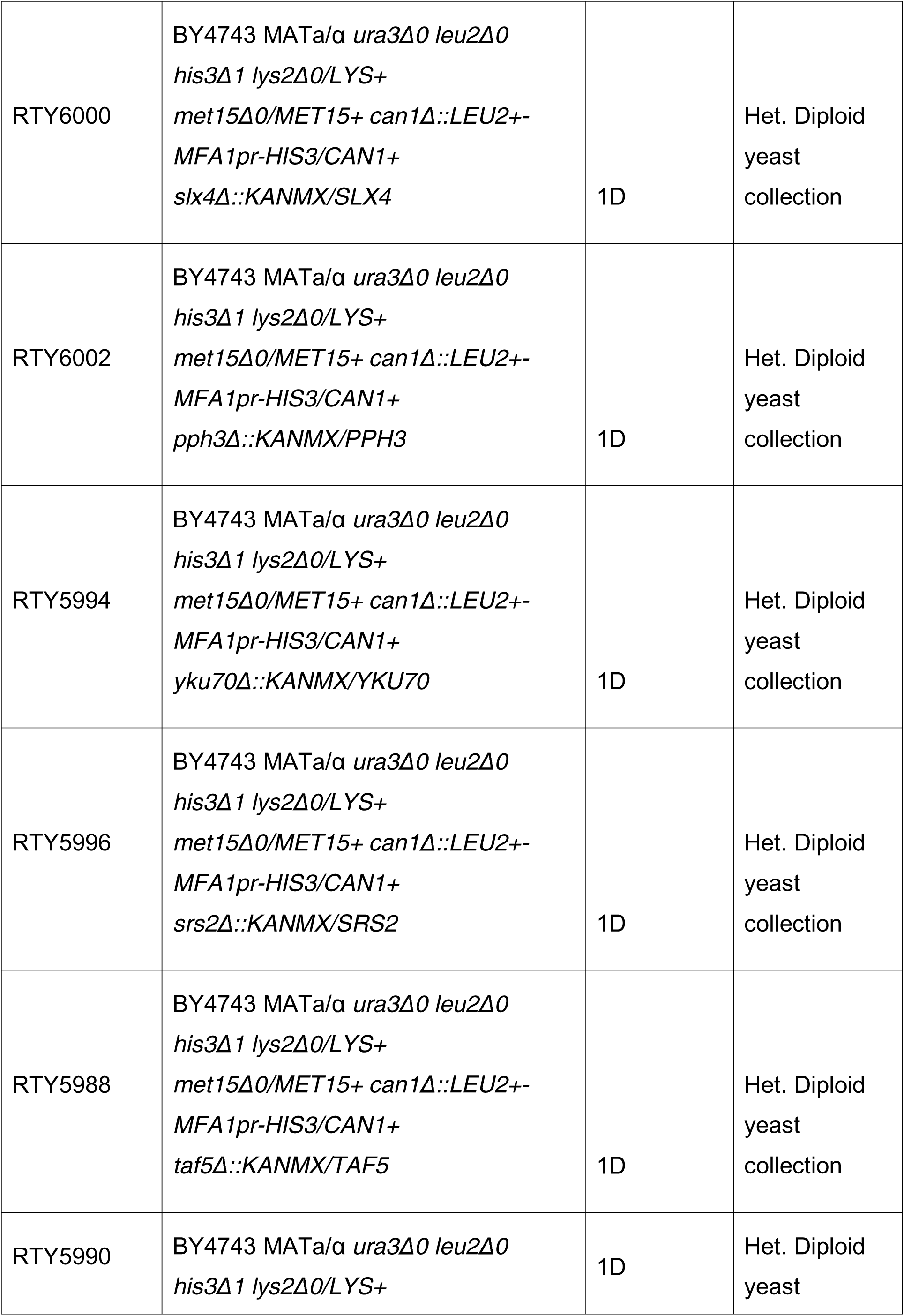
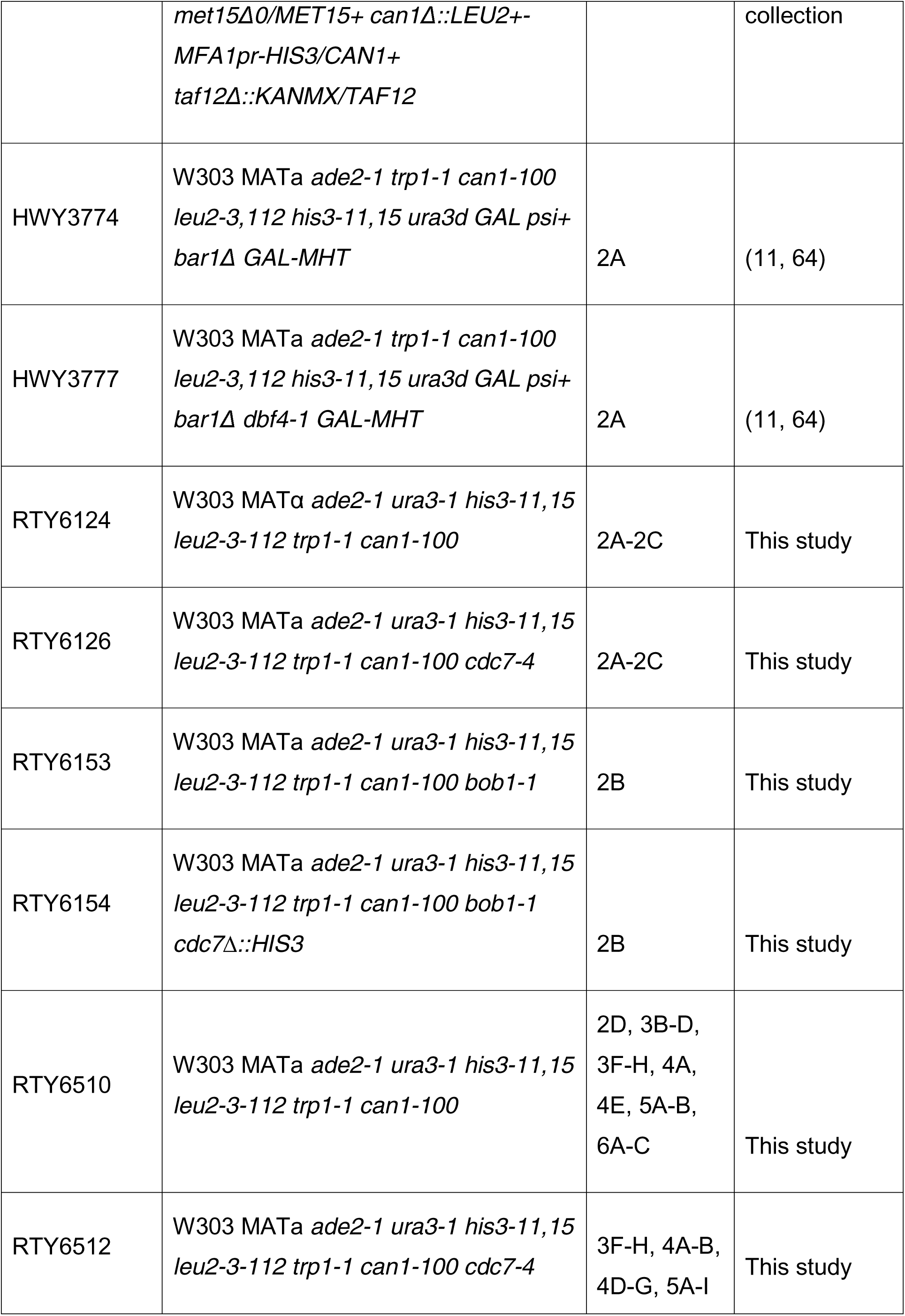
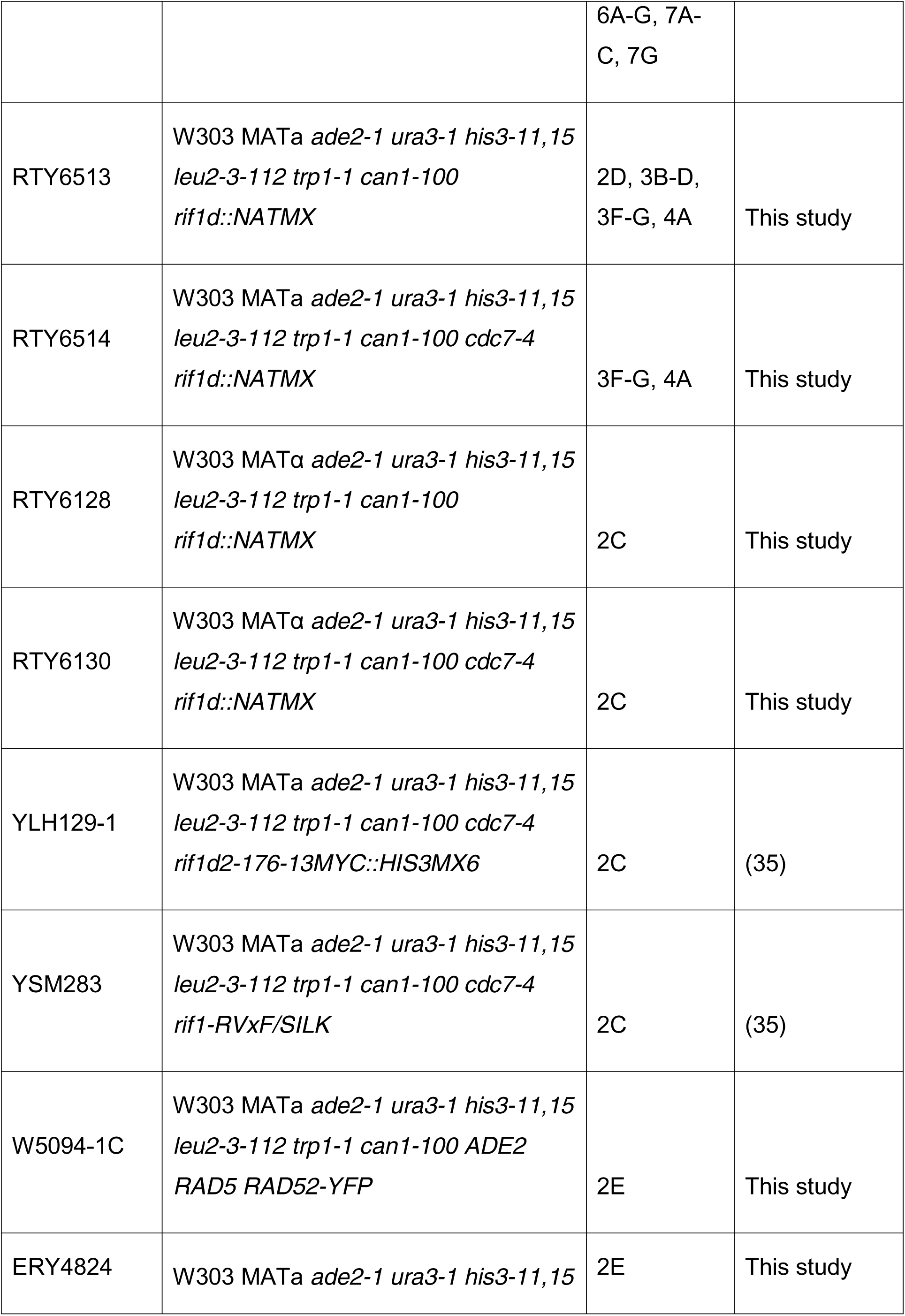
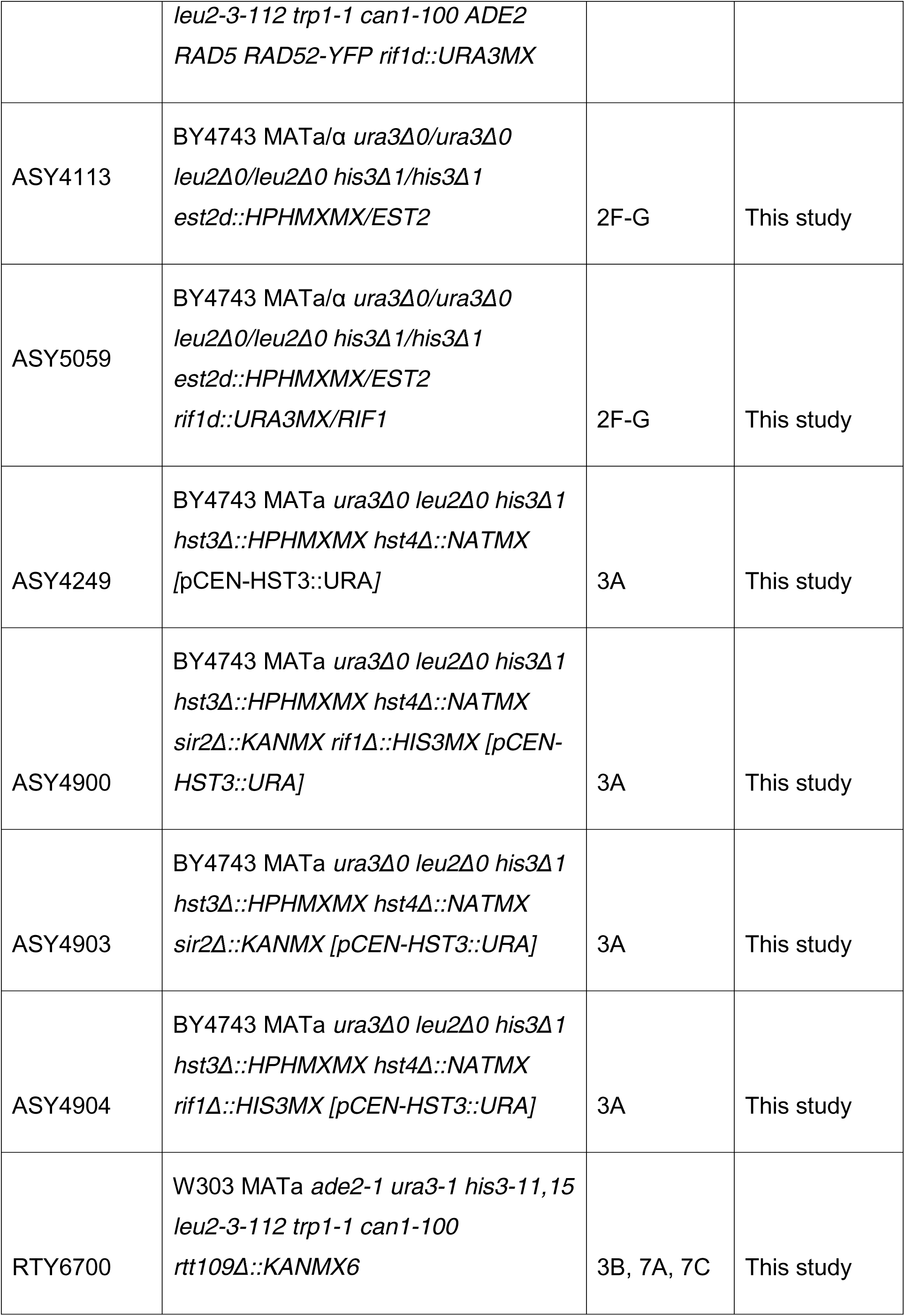
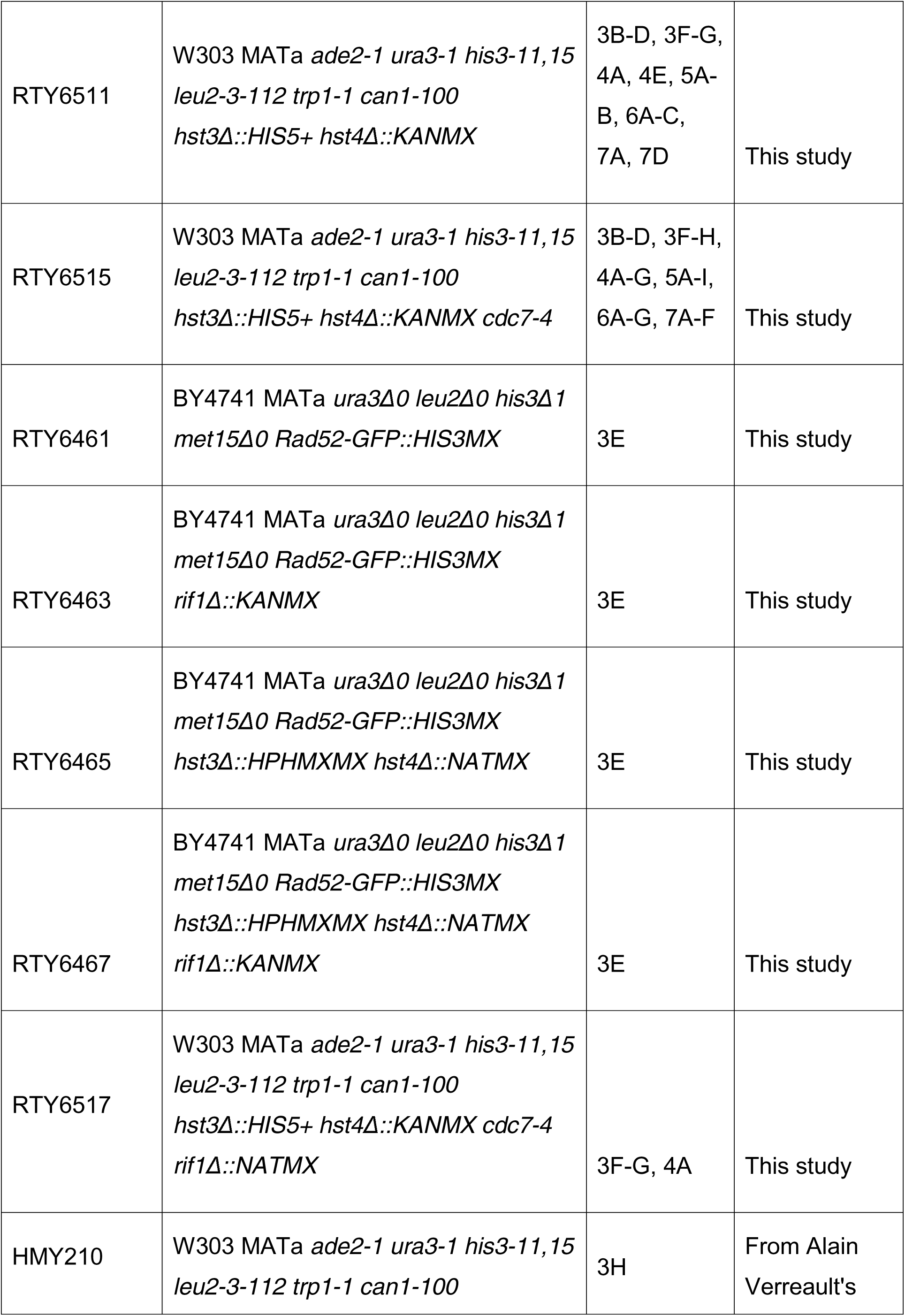
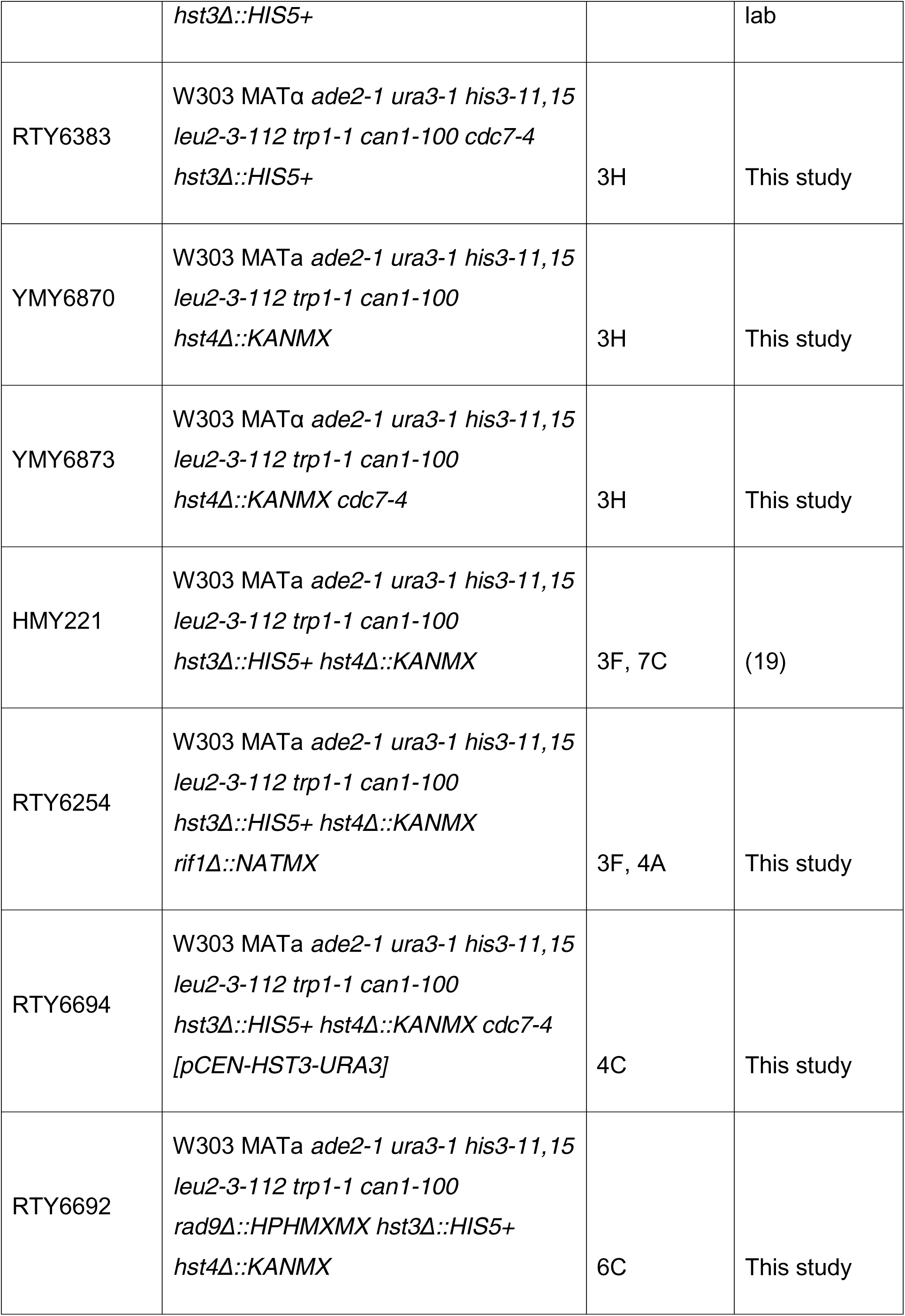
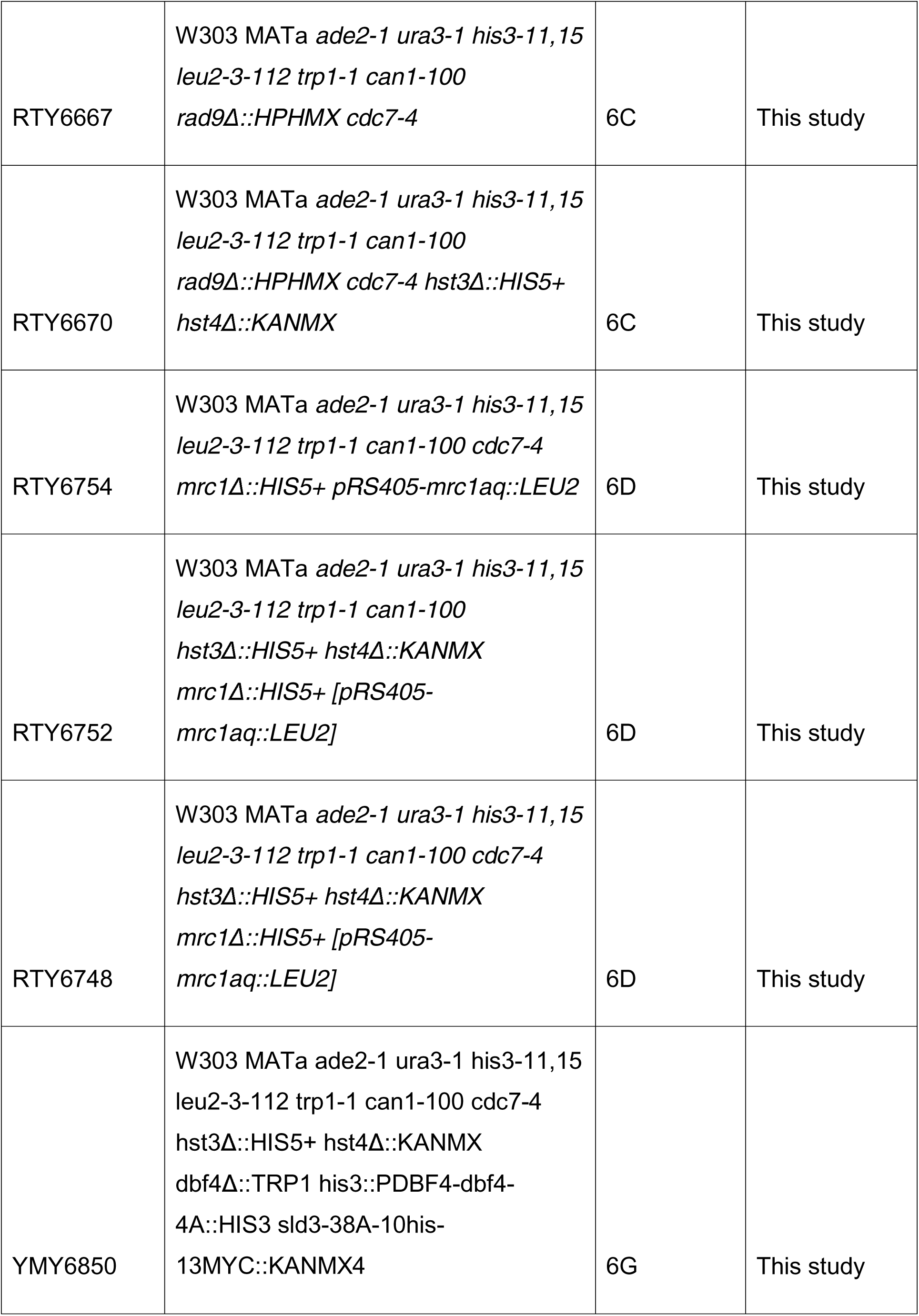
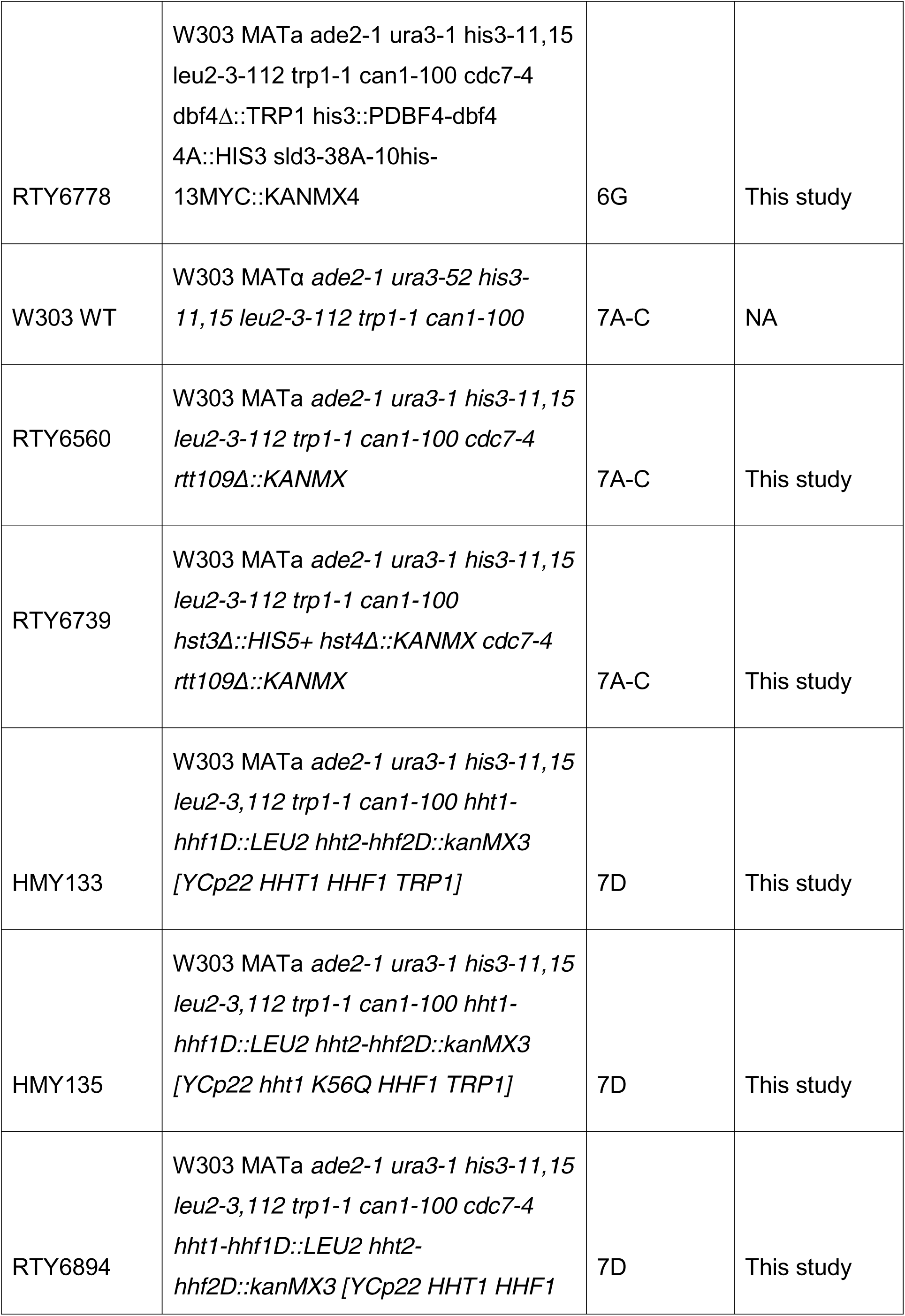
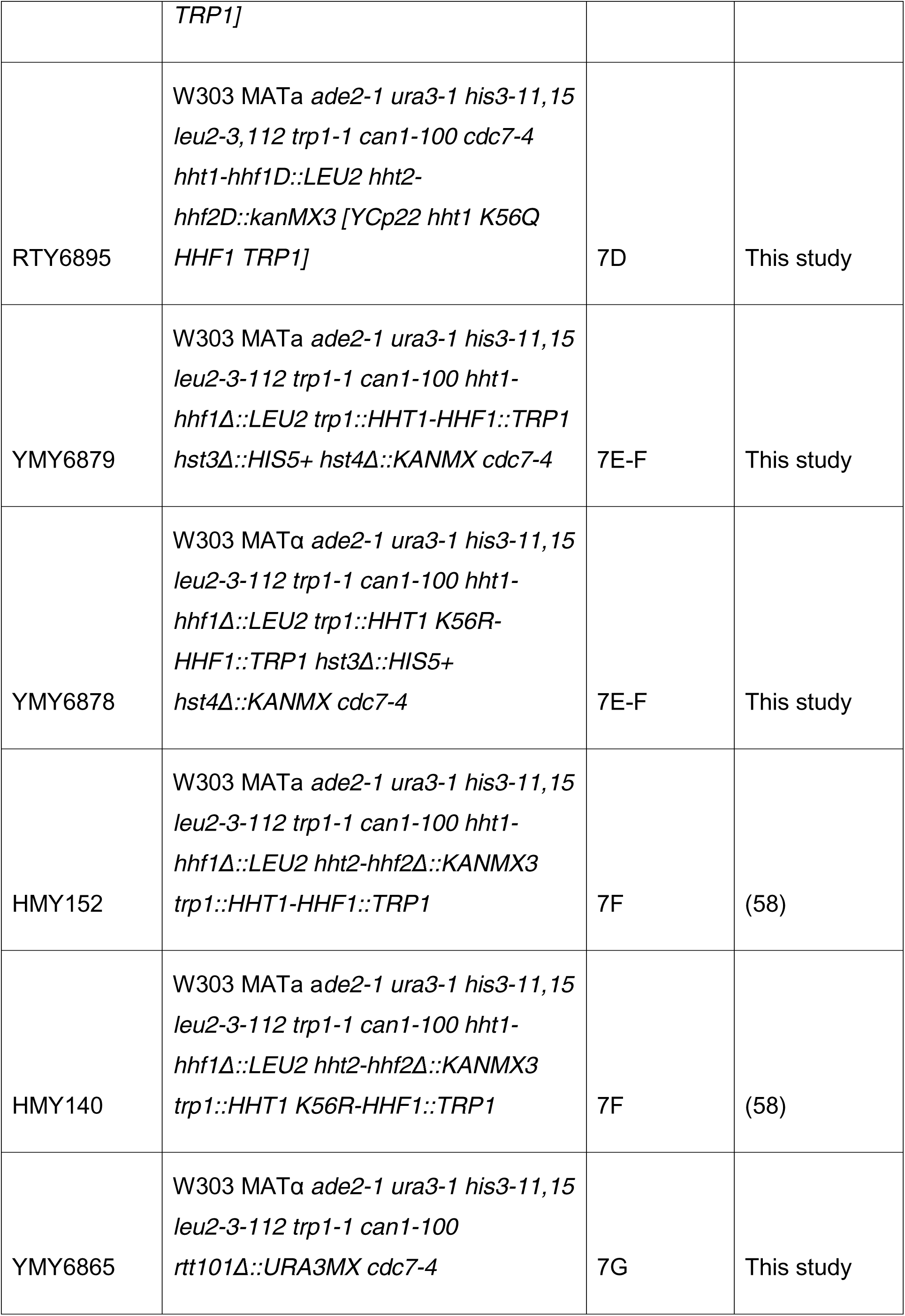
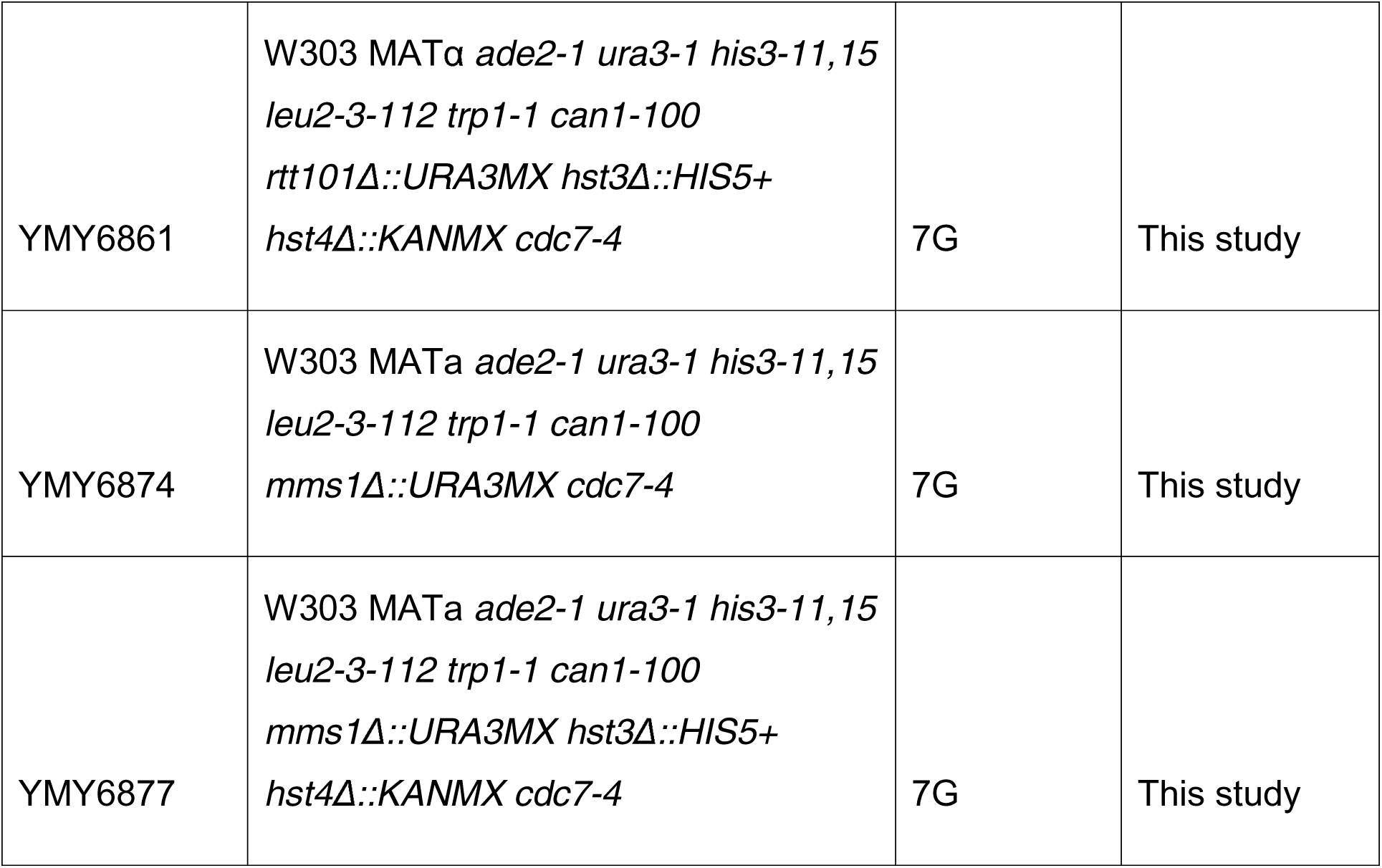
Yeast strains used in this study

**TABLE 2:**
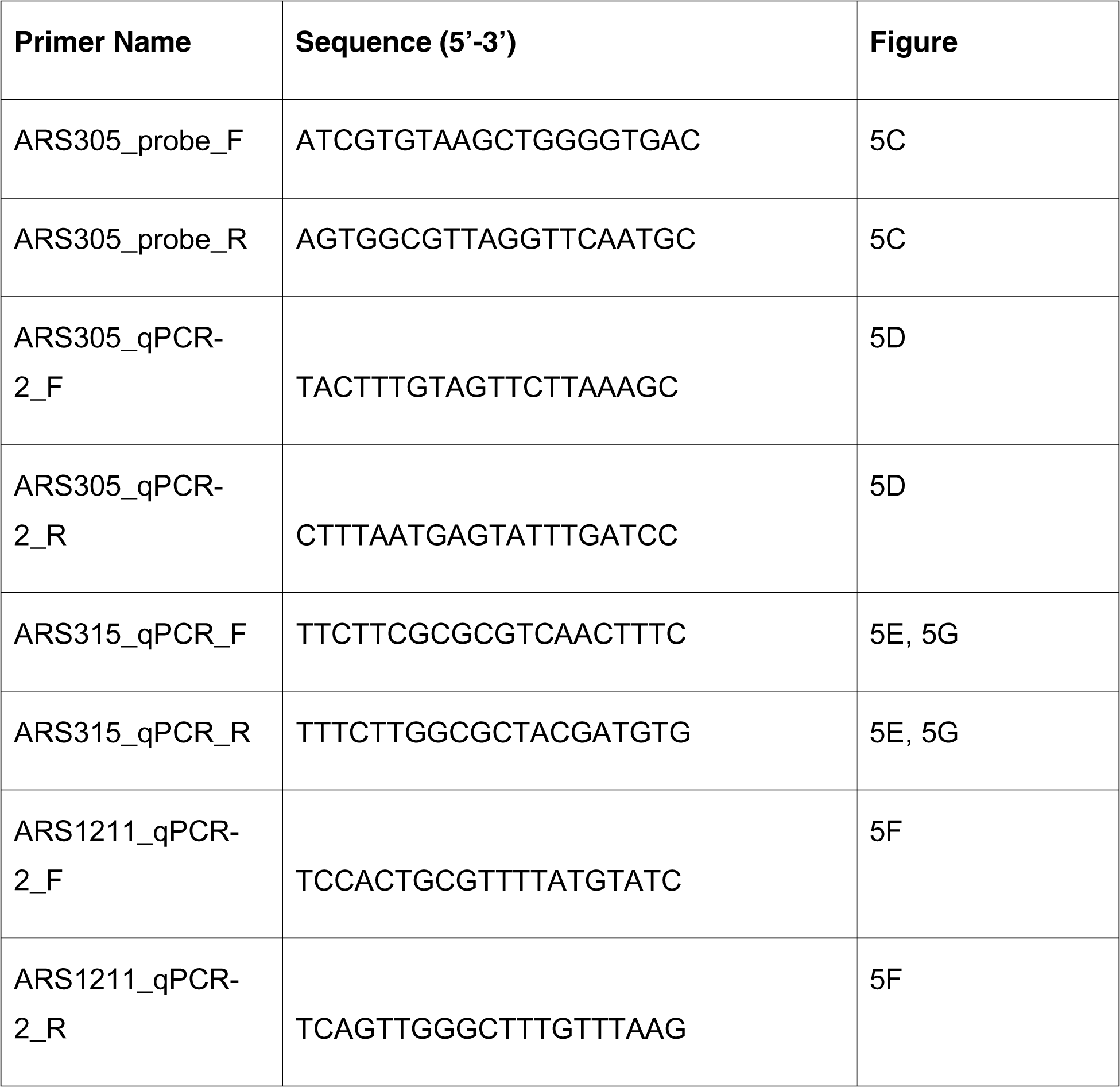
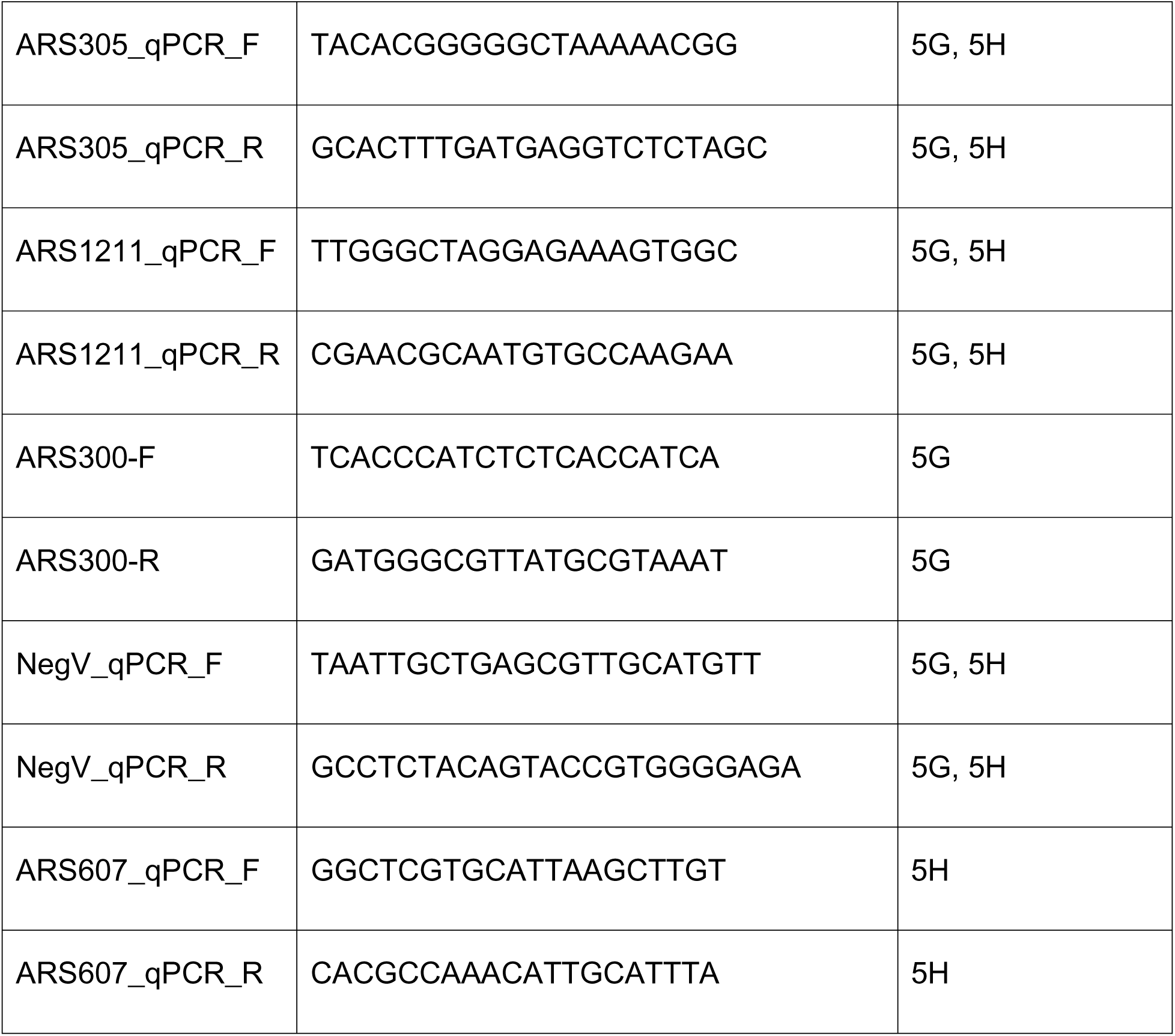
PCR primers used in this study.

We previously demonstrated that transient exposure to methyl methane sulfonate (MMS), an alkylating agent that generates replication-blocking lesions such as 3-methyl adenine, prevents timely completion of S phase in *hst3*Δ *hst4*Δ cells (24). DNA content flow cytometry analyses revealed that deletion of *RIF1* noticeably rescued the S phase progression delay caused by transient MMS exposure in *hst3*Δ *hst4*Δ double mutants (Figure 3C), consistent with the notion that Rif1 compromises DNA replication completion in these cells. Nevertheless, deletion of *RIF1* did not rescue the sensitivity of *hst3*Δ *hst4*Δ to MMS (Figure 3D), which might be due to the fact that Rif acts to stabilize stalled RF (41) in addition to its role in regulating origin activity. We also found that *rif1*Δ reduced spontaneous formation of Rad52 foci and histone H2A S129 phosphorylation in *hst3*Δ *hst4*Δ cells (Figure 3E-F), indicating that, in the absence of exogenous replicative stress-inducing genotoxins, Rif1 activity causes DNA damage in Hst3/Hst4-deficient cells. The above data, combined with those linking Rif1 to NAM sensitivity, support the notion that Rif1/Glc7-mediated reversal of DDK-dependent phosphorylation, and consequent inhibition of origins of DNA replication, contributes to the phenotypes of cells lacking Hst3 and Hst4. Consistently, deletion of *HST3* and *HST4* considerably exacerbated the temperature sensitivity of *cdc7-4* mutant cells in a Rif1-dependent manner (Figure 3G). Deletion of either *HST3* or *HST4* alone did not increase the temperature sensitivity of *cdc7-4* (Figure 3H), in accord with the known functional redundancy of Hst3/Hst4 with respect to histone deacetylation (19, 22). Altogether, these results suggest that reduction in replication origin activity is detrimental to cells lacking Hst3/Hst4.

### *cdc7-4 hst3*Δ *hst4*Δ cells display synthetic defects in the initiation of origins of replication

We sought to further explore DNA replication dynamics in *cdc7-4* cells lacking Hst3/Hst4. To this end, we synchronized *cdc7-4 hst3*Δ *hst4*Δ and appropriate control cells in G1 at the permissive temperature of 25°C using alpha factor, followed by release toward S phase at the semi-permissive temperature (for *cdc7-4*) of 25°C. Strikingly, we found that *cdc7-4 hst3*Δ *hst4*Δ cells displayed strong inhibition of S phase progression when released from G1 toward S at 30°C compared to either *hst3*Δ *hst4*Δ or *cdc7-4* (Figure 4A). Such S phase progression defect was not observed at the permissive temperature of 25°C, indicating that the impact of reduced Cdc7 activity (due to incubation at the semi-permissive temperature of 30°C for *cdc7-4*) on DNA replication is strongly exacerbated by deletion of *HST3* and *HST4* (Figure 4B). As expected, this phenotype was rescued by deletion of *RIF1* (Figure 4A) or expression of plasmid-borne copy of *HST3* (Figure 4C). The observed DNA replication defect does not appear to result from compromised release from alpha factor-mediated G1 arrest, since asynchronously growing *cdc7-4 hst3*Δ *hst4*Δ cells were also found to accumulate in early S when incubated at 30°C (Figure 4D). Moreover, the budding index of *cdc7-4 hst3*Δ *hst4*Δ cells released from alpha factor-mediated G1 block toward S phase at 30°C was comparable to that of *cdc7-4* cells (≈ 50-60% of cells with detectable buds; Figure 4E) even though the former cells present barely detectable S phase progression in these conditions (Figure 4A). We note that the small size of buds at 45 and 60 minutes post-release from alpha factor rendered precise assessment of budding index challenging. To further confirm our results, we performed an experiment in which *cdc7-4* and *cdc7-4 hst3*Δ *hst4*Δ cells were released from alpha factor-induced G1 arrest toward S at the non-permissive temperature of 39°C for 3h, thereby allowing time for buds to become larger while preventing Cdc7 activity and, consequently, initiation of DNA synthesis at origins (Figure 4F). After monitoring the budding index, the temperature of the culture was decreased to 30°C for 30 minutes to evaluate S phase progression (Figure 4F). While for unknown reasons the fraction of *cdc7-4 hst3*Δ *hst4*Δ *and cdc7-4* mutants with detectable buds did not reach more than 60 to 80%, respectively, in these conditions (Figure 4G), S phase progression remained completely blocked in *cdc7-4 hst3*Δ *hst4*Δ, but not *cdc7-4* cells, after incubation at 30°C (Figure 4F). Overall, the data indicate that even though *cdc7-4 hst3*Δ *hst4*Δ cells enter S phase at the semi-permissive temperature of 30°C, DNA replication progression is strongly inhibited in these conditions.

**FIGURE 4.**
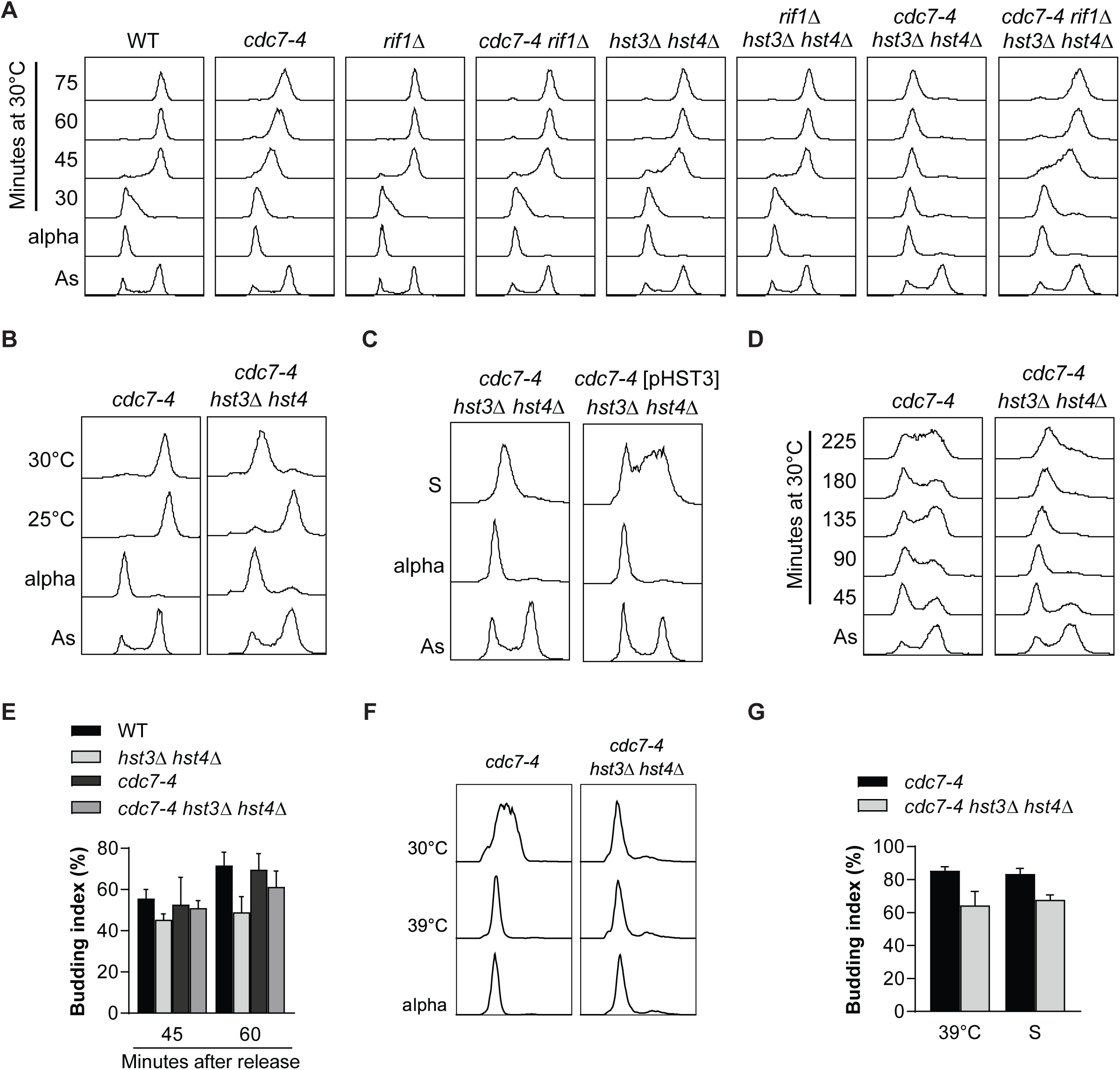
Deletion of *HST3* and *HST4* inhibits S phase progression in *cdc7-4* cells. (A) Cells were arrested in G1 using alpha factor at 25°C (alpha) and released toward S phase at 30°C. Samples were taken for DNA content analysis by flow cytometry at the indicated time points. As: Asynchronous. (B) Cells were arrested in G1 using alpha factor at 25°C (alpha) and released toward S phase at 25°C or 30°C for 90 minutes before harvest. Samples were taken for DNA content analysis by flow cytometry. As: Asynchronous. (C) Cells were treated as in A. Samples were collected 90 minutes after release from alpha factor at 30°C. (D) Exponentially growing cells at 25°C were transferred to 30°C for the indicated time and harvested for DNA content analysis by flow cytometry. As: Asynchronous. (E) Budding index was assessed 45 and 60 minutes after release from alpha-factor toward S at 30°C. Cells were treated as in A. At least 100 cells were inspected per condition. (F) Cells were arrested in G1 at 25°C using alpha factor (alpha) and released toward S at 39°C for 3 h. Cells were then transferred to 30°C for 30 minutes before harvest. Samples were taken for DNA content analysis by flow cytometry. (G) Budding index of cells harvested in D. At least 100 cells were inspected per condition.

Given the known role of Dbf4-Cdc7 in activating MCM helicase complexes at origins, we suspected that cdc7-4 *hst3*Δ *hst4*Δ cells might present synthetic defects in the initiation of DNA replication. Formation of RF at origins prevents the entry of yeast chromosomes in Pulsed-Field Gel Electrophoresis (PFGE) gels (42). We found that PFGE signals, reflecting entry of chromosomes in the gel, were significantly stronger in *cdc7-4 hst3*Δ *hst4*Δ cells at 45 and 60 minutes after release from alpha factor compared to WT, *hst3*Δ *hst4*Δ and *cdc7-4* (Figure 5A-B). This is consistent with the notion that a reduced proportion of cells activated origins throughout chromosomes in *cdc7-4 hst3*Δ *hst4*Δ cells compared to control strains. We next used alkaline gel electrophoresis and Southern blotting to detect formation of low molecular weight nascent DNA at the efficient early origin ARS305, as described in (6). The results indicate a strong reduction in the amount of low molecular weight DNA formed at this origin within 60 minutes of release from alpha factor arrest at 30°C in *cdc7-4 hst3*Δ *hst4*Δ cells compared to *cdc7-4* mutants (Figure 5C).

**FIGURE 5.**
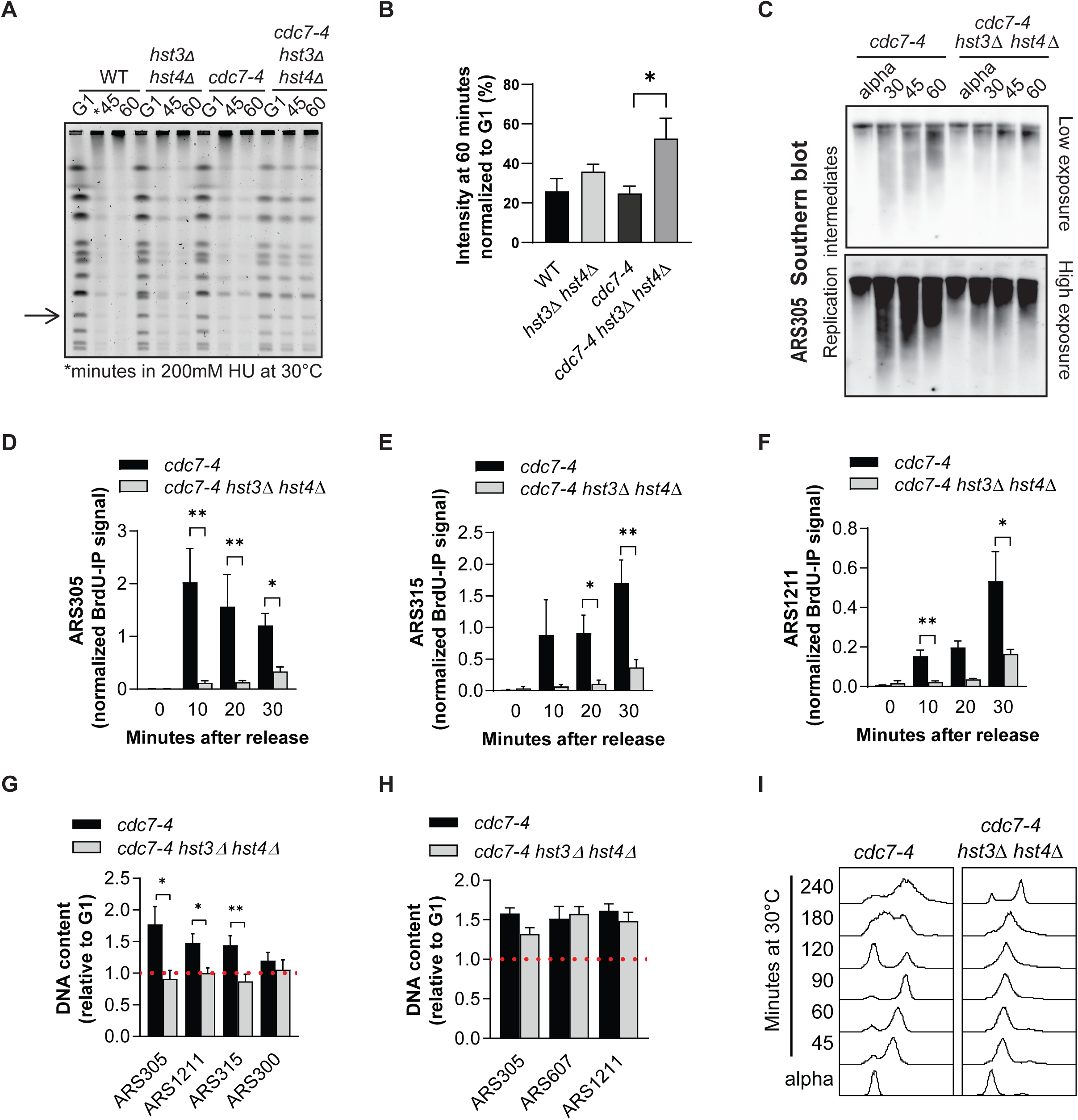
Deletion of *HST3* and *HST4* inhibits the activation of origins of replication in *cdc7-4* cells. (A) Cells were arrested in G1 at 25°C using alpha factor (G1) and released toward S phase in presence of 200 mM HU for the indicated time at 30°C. PFGE was performed as described in Material and Methods. The arrow indicates the band used for quantification in B. (B) Densitometric quantification of the selected band from A. Bars represent the mean ± SEM of 3 independent experiments. (C) Cells were arrested in G1 using alpha factor at 25°C (alpha) and released at 37°C for 1 h. Cells were then transferred to 30°C for the indicated time period before harvest. DNA samples were run on alkaline gels followed by Southern blotting to detect short ssDNA fragments generated at ARS305 during origin activation. (D-F) Cells were arrested in G1 at 25°C using alpha factor and released at 37°C for 1 h. Cells were then transferred to 30°C for 30 minutes, or to 25°C for 1h in presence of 200 mM HU before harvest (for normalization purposes, see Material and Methods). DNA samples were extracted, immunoprecipitated using anti-BrdU antibody and processed for quantitative PCR analysis. Bars represent the mean ± SEM of the percent of the input of 5 independent experiments using qPCR primers for the early origins (D) ARS305, (E) ARS315 and (F) ARS1211. Normalization for BrdU intake capacity per strain was performed using the 25°C for 1h in presence of 200 mM HU as described in Material and Methods. (G) Cells were arrested in G1 using alpha factor at 25°C and released toward S at 37°C for 1 h. Cells were then incubated at 30°C for 30 minutes before harvest. DNA was extracted and processed for quantitative PCR analysis (see Methods). qPCR signal for a given origin was normalized to that obtained from the NegV locus (which is expected to be unreplicated 30 minutes post-release from G1 toward S), and then divided by the normalized signals obtained from alpha-factor arrested (G1) cells. Graph bars represent mean ± SEM of five independent experiments. (H) qPCR analysis of DNA content at selected origins. Indicated strains were treated as in G, except that cells were released toward S phase in presence of 200 mM HU for 120 minutes at 30°C before harvest. Graph bars represent mean ± SEM of three independent experiments. (I) Cells were arrested in G1 using alpha factor (alpha) at 25°C and released toward S phase at 30°C for the indicated time. Samples were taken for DNA content analysis by flow cytometry. As: asynchronous. Throughout this figure, *: p < 0.05 and **: p < 0.01, unpaired two-tailed Student’s t-test.

To further compare origin initiations in *cdc7-4 hst3*Δ *hst4*Δ vs *cdc7-4* cells, we released G1-arrested cells toward S phase at the non-permissive temperature of 37°C in the presence of the nucleoside analog BrdU for 60 minutes, and then switched the temperature of the cultures to 30°C for 30 minutes. BrdU-IP followed by quantitative PCR (qPCR) was then used to quantify incorporation of BrdU in genomic DNA at three early origins (ARS305, ARS315 and ARS1211). This analysis revealed that BrdU incorporation into nascent DNA is significantly reduced in *cdc7-4 hst3*Δ *hst4*Δ compared to *cdc7-4* cells at these early/efficient origins of replication (Figure 5D-F). Consistently, qPCR analysis on total genomic DNA showed that compared to *cdc7-4* cells, duplication of DNA at these origins was inhibited in *cdc7-4 hst3*Δ *hst4*Δ mutants 30 minutes after release from alpha factor arrest at 30°C (Figure 4G). We note that *cdc7-4 hst3*Δ *hst4*Δ cells eventually initiated DNA replication and completed S phase when incubated for extended periods at 30°C (240 minutes post-release from alpha factor arrest; Figure 4G-H). Taken together, the results indicate that the *hst3*Δ *hst4*Δ mutations cause synthetic defects in the activation of replication origins when combined with *cdc7-4*, thereby strongly delaying S phase progression at 30°C.

### Inhibition of origin activity in *cdc7-4 hst3***Δ** *hst4***Δ** cells is not due to activation of Rad53 in early S phase

One of the key roles of intra-S phase checkpoint signaling is to limit the activation of origins in response to replication stress (6, 43). In yeast, this has been shown to occur via Rad53-dependent phosphorylation and consequent inactivation of Dbf4 and Sld3 (7). Cells lacking Hst3/Hst4 activity are known to present spontaneous DNA damage and constitutive activation of Rad53 (19, 23, 24, 26, 27). We therefore sought to investigate the influence of Rad53 activity on the S phase progression delay observed in *cdc7-4 hst3*Δ *hst4*Δ cells. While *cdc7-4 hst3*Δ *hst4*Δ cells constitutively present some Rad53 phosphorylation in G1, no obvious elevation in Rad53 autophosphorylation-induced electrophoretic mobility shift was observed upon release of cells from alpha factor arrest toward S phase at 30°C, either in the presence or absence of the replication-blocking drug hydroxyurea (HU; Figure 6A-B). This result is consistent with the notion that few if any RF are progressing in these conditions (Figure 5D-G), thereby reducing the number of stalled RF in either HU-treated or untreated conditions and consequent Rad53 activation.

**FIGURE 6.**
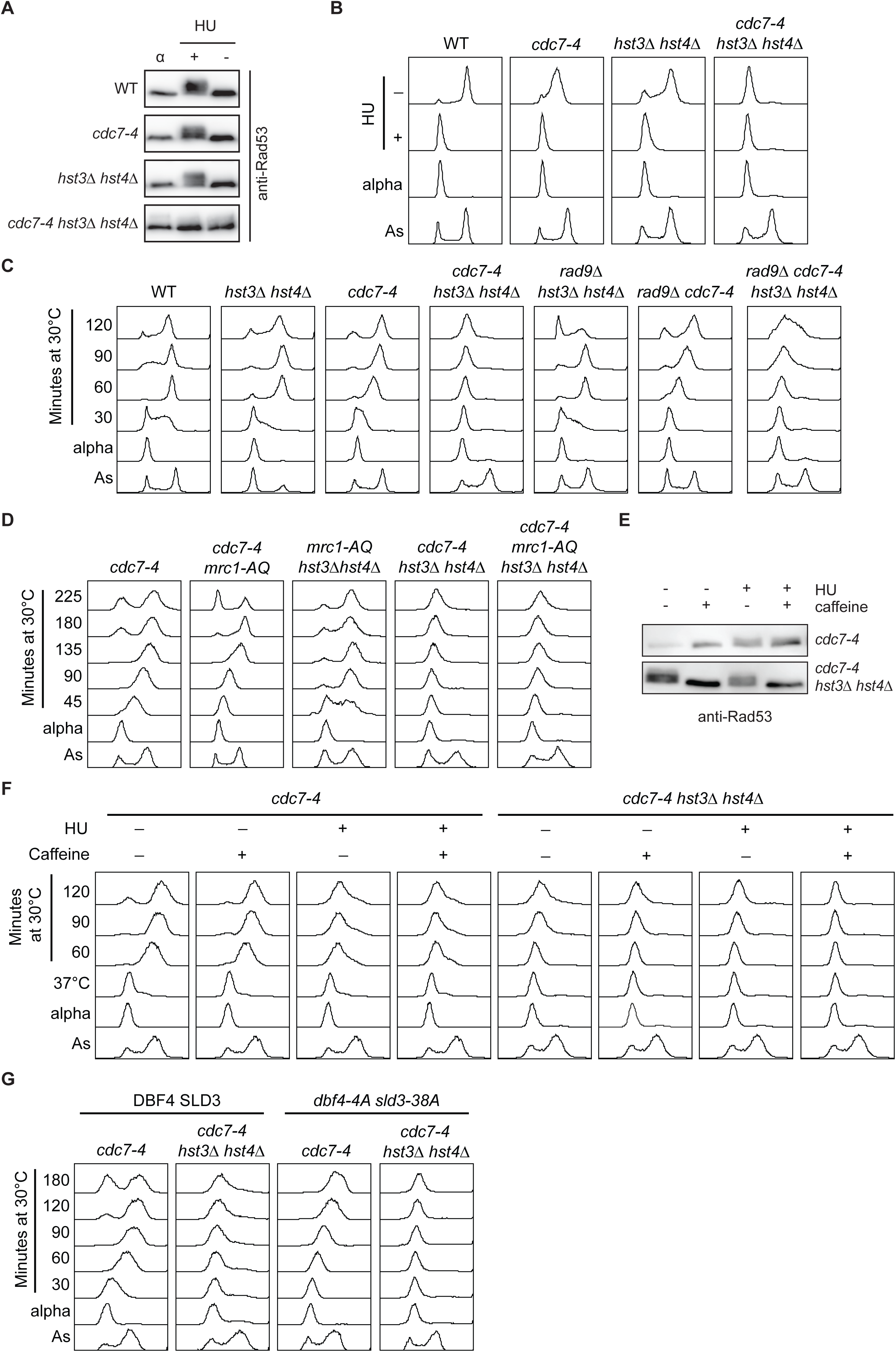
The DNA replication defects of *cdc7-4 hst3*Δ *hst4*Δ cells are not due to Rad53 activation in early S phase. (A-B) Cells were arrested in G1 at 25°C using alpha factor (alpha) and released toward S phase at 30°C in the presence or absence of 200 mM HU. Cells were harvested 60 minutes post-release toward S and processed for immunoblotting (A) and DNA content analysis by flow cytometry (B). As: Asynchronous. (C) Cells were arrested in G1 at 25°C using alpha factor (alpha) and released toward S phase at 30°C. Samples were taken for DNA content analysis by flow cytometry. As: Asynchronous. (D) DNA content analysis by flow cytometry. Cells were treated as in C. (E-F) Cells were treated as in C except that releases toward S phase at 30°C were done in YPD +/- 200 mM HU +/- 0.15% caffeine. After 60 minutes of release, cells were harvested for immunoblotting (E). DNA content was assessed by flow cytometry using samples harvested at the indicated time points (F). (G) DNA content analysis by flow cytometry. Cells were treated as in C.

We next tried to delete *RAD53* and *SML1* in *cdc7-4 hst3*Δ *hst4*Δ cells to directly assess the role of intra-S phase checkpoint signaling on the phenotypes of these mutants; *SML1* deletion is necessary to permit viability of *rad53*Δ mutants (44). However, even though *hst3*Δ *hst4*Δ *rad53*Δ *sml1*Δ cells are viable (23), we failed to generate a *cdc7-4 hst3*Δ *hst4*Δ *rad53*Δ *sml1*Δ, suggesting that for unknown reason this combination of mutations causes synthetic lethality. To circumvent this, we engineered *cdc7-4 hst3*Δ *hst4*Δ strains harboring mutations in *MRC1* and *RAD9*, two key mediators of the activation of the intra-S phase checkpoint (5). We found that deletion of *RAD9* in *cdc7-4 hst3*Δ *hst4*Δ cells led to modest improvement in S phase progression, but only after 60 minutes of release from alpha factor arrest (Figure 6C). This suggests that Rad9 might contribute to the long-term maintenance, rather than the establishment, of S phase progression defects in *cdc7-4 hst3*Δ *hst4*Δ cells. In contrast, expression of a mutated allele of Mrc1 (*mrc1-AQ*) which compromises its role in activating intra-S phase checkpoint kinases (45) did not have any influence on S phase progression in *cdc7-4 hst3*Δ *hst4*Δ cells (Figure 6D). We further found that while inhibition of the apical kinase of the intra-S phase checkpoint Mec1 using caffeine (46, 47) completely abrogated Rad53 phosphorylation, as expected, such treatment did not rescue the strong inhibition of DNA replication progression of *cdc7-4 hst3*Δ *hst4*Δ mutants (Figure 6E-F). We also note that caffeine did not prevent *cdc7-4* cells from completing DNA replication at 30°C (Figure 6F). Expression of Dbf4 and Sld3 variants that cannot be phosphorylated by Rad53 was previously shown to abrogate intra-S phase checkpoint-dependent inhibition of origin activity in yeast (7). We found that introducing such mutated alleles of *DBF4* and *SLD3* in *cdc7-4 hst3*Δ *hst4*Δ cells does not alleviate their S phase progression defects at 30°C; moreover, mutations of *DBF4* and *SLD3* did not prevent *cdc7-4* cells from completing S phase in these conditions (Figure 6G). We conclude that the incapacity of *cdc7-4 hst3*Δ *hst4*Δ mutants to initiate DNA replication in a timely manner at the beginning of S phase when released from G1 at 30°C is not due to Rad53-dependent phosphorylation of Sld3 and Dbf4 and consequent inhibition of early origins of replication.

### Constitutive histone H3 lysine 56 acetylation causes replication defects in *cdc7-4* cells

Constitutive acetylation of histone H3 lysine 56 (H3 K56ac) causes most of the phenotypes associated with *hst3*Δ *hst4*Δ mutants, including their temperature and DNA damage sensitivity (19, 22, 23). While H3 K56ac strictly depends on the Rtt109 histone acetyltransferase (48), this acetyltransferase also acetylates other residues in the N-terminal tail of histone H3 (21, 49, 50), although there are currently no evidence that link the acetylation of these residues with the phenotypes of *hst3*Δ *hst4*Δ mutants. We found that deletion of *RTT109* significantly rescued DNA replication progression and growth of *cdc7-4 hst3*Δ *hst4*Δ cells at semi-permissive temperatures for *cdc7-4* (between 28°C and 30°C; Figure 7A-C). Moreover, replacing histone H3 by a H3 K56Q variant to mimic constitutive H3 K56ac in *cdc7-4* cells caused strong synthetic temperature sensitivity (Figure 7D).

**FIGURE 7.**
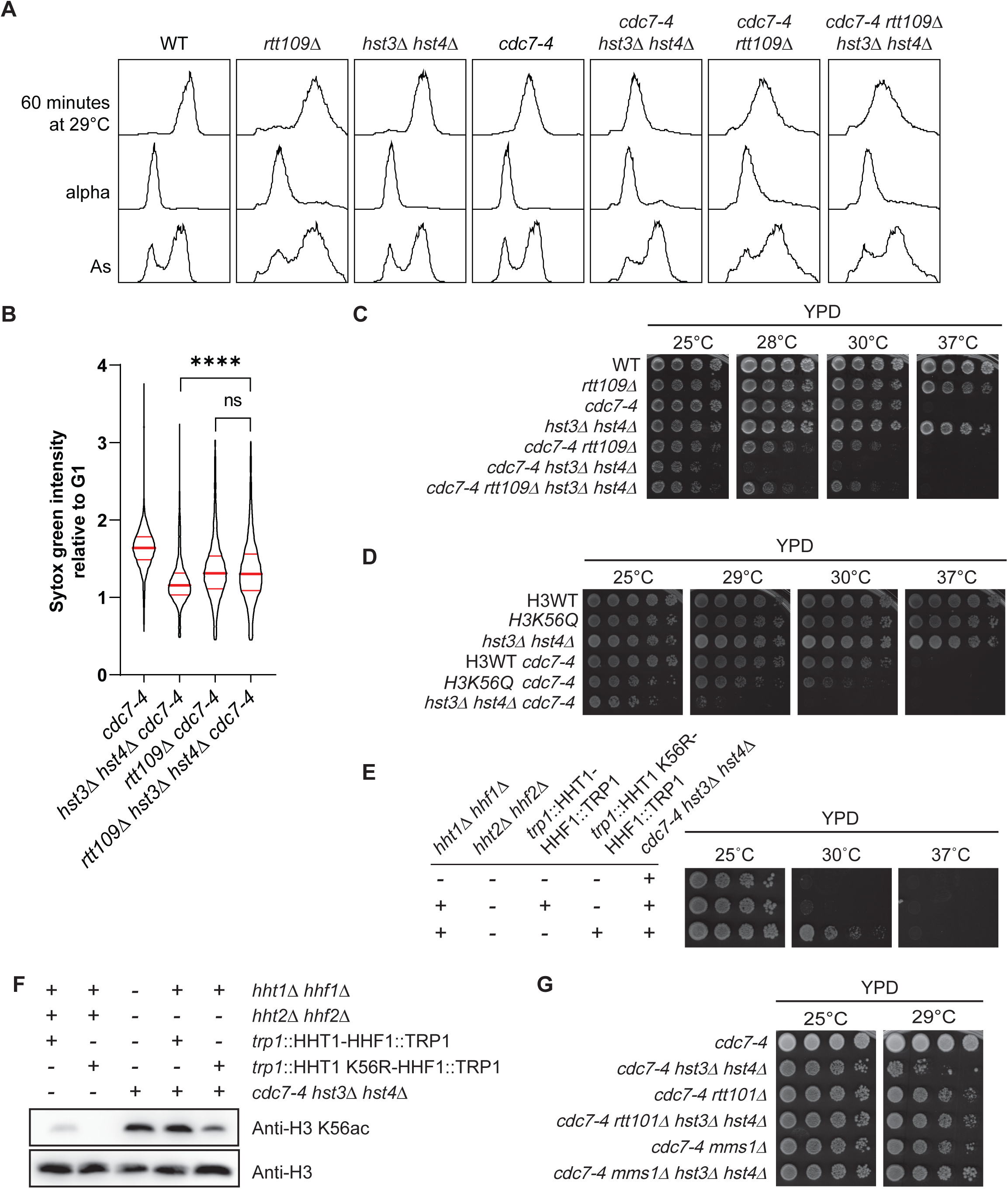
Constitutive H3K56 acetylation, Rtt101, and Mms1 cause the S phase progression defects and synthetic temperature sensitivity of *cdc7-4 hst3*Δ *hst4*Δ cells. (A-B) Cells were arrested in G1 at 25°C using alpha factor (alpha) and released toward S for 60 minutes at 29°C. Samples were taken for DNA content analysis by flow cytometry. (B) Violin plot represents the Sytox Green value (DNA content) per cell from the 60 minutes time point in A normalized to their corresponding G1 median value. Red bars represent the median and quartiles. ns: p value > 0.05 and ****: p value < 0.0001, unpaired two-tailed Mann-Whitney test. (C) 5-fold serial dilutions of cell cultures were spotted on YPD-agar plates. Plates were incubated at the indicated temperature. (D) As in C. (E) As in C. (F) Exponentially growing cells at 25°C were processed for immunoblotting. (G) As in C.

We next sought to engineer a *cdc7-4 hst3*Δ *hst4*Δ strain lacking H3 K56ac via expression of a histone H3 variant in which lysine 56 is replaced by a non-acetylable arginine residue (H3 K56R). To this end, both copies of the endogenous genes encoding histone H3 (*HHT1* and *HHT2*) were deleted while one copy of the *HHT1* gene +/- K56R mutation was integrated at the *TRP1* locus. We failed to generate the *cdc7-4 hst3*Δ *hst4*Δ cells expressing either H3 WT or H3 K56R using this standard strategy, suggesting that abnormal histone gene dosage due to deletion of *HHT2* may be lethal in this context. To circumvent this issue and reduce H3 K56ac levels without changing histone gene dosage, we replaced *HHT1* by either a WT or K56R allele and left the endogenous copy of *HHT2* intact (Figure 7E-F). This method produced viable *cdc7-4 hst3*Δ *hst4*Δ strains in which H3 K56ac levels are either unchanged (H3 WT) or noticeably reduced (H3 K56R; Figure 7E-F). Strikingly, we observed a strong rescue of the temperature sensitivity of *cdc7-4 hst3*Δ *hst4*Δ upon expression of H3 K56R (compared to control cells expressing H3 WT; Figure 7E). Overall, the above results indicate that constitutive Rtt109-dependent H3 K56ac underlies the synthetic temperature sensitivity of *cdc7-4 hst3*Δ *hst4*Δ cells.

Genetic and biochemical data indicate that Rtt109 and H3 K56ac act at least in part by modulating the activity of a ubiquitin ligase complex composed of the Rtt101, Mms1 and Mms22 subunits (51–54). Deletion of the genes encoding subunits of this complex partially suppresses the phenotypes of *hst3*Δ *hst4*Δ cells (51), although the precise mechanisms linking constitutive acetylation of nucleosomal H3 K56ac with Rtt101/Mms1/Mms22 is incompletely characterized. We found that deletion of either *RTT101* or *MMS1* suppressed the synthetic temperature sensitivity of *cdc7-4 hst3*Δ *hst4*Δ mutant cells (Figure 7G). In contrast, we were unable to generate *cdc7-4 hst3*Δ *hst4*Δ *mms22*Δ cells, suggesting that synthetic lethal interactions between these genes prevent viability. Nevertheless, our results implicate Rtt101-Mms1-containing complexes in H3 K56ac-dependent modulation of DNA replication origins.

## DISCUSSION

In yeast, virtually all new histones H3 are acetylated on K56, leading to a chromosome-wide wave of H3 K56ac during S phase (19, 48). This modification promotes timely formation of nucleosomes behind RF by favoring the interaction of new histones with the chromatin assembly factors CAF1 and Rtt106 (55). While this “pre-deposition” function of H3 K56ac is well-established, several observations suggest that this mark also plays important biological roles following its incorporation into chromatin (56). Constitutive nucleosomal H3 K56ac causes spontaneous DNA damage and extreme sensitivity to replication-blocking drugs in *hst3*Δ *hst4*Δ cells, suggesting that chromatin-associated H3 K56ac influences the cellular response to replicative stress (22, 23). Moreover, cells have evolved molecular mechanisms to degrade Hst3 in response to replicative stress (57, 58), which raises the possibility that the ensuing persistence of H3 K56ac may somehow contribute to the DNA damage response. Nevertheless, while a multitude of cellular pathways have been associated with nucleosomal H3 K56ac (23, 24, 26, 27, 51), the molecular basis of the sensitivity of *hst3*Δ *hst4*Δ mutants to replicative stress, as well as the role of H3 K56ac persistence after DNA damage, are poorly understood.

We previously showed that NAM-induced inhibition of Hst3 and Hst4 causes replicative stress by elevating H3 K56ac (26, 27). In accord with this, our current and previously published screens (26) revealed that several genes conferring NAM resistance participate in DNA replication and repair (Figure 1C-D). Interestingly, the current screen also revealed that lack of *TAF5* and *TAF12*, which encodes proteins shared by TFIID and the SAGA acetyltransferase complex, strongly sensitizes cells to NAM. Since the SAGA complex modulates the expression of stress-responsive genes (59), we speculate that the presence of subunits of this complex among the “hits” of our screen might reflect transcriptional activation of critical cellular stress responses pathways during NAM treatment. Our screen also identified genes involved in proteasome regulation and ubiquitin-dependent processes as modulators of NAM sensitivity (Figure 1C). As mentioned previously, the Rtt101-Mms1-Mms22 ubiquitin ligase complex displays clear genetic links with H3 K56ac in the context of the response to replicative stress (51–54). It is therefore possible that heterozygosity in genes involved in ubiquitin- and proteasome-related processes elevate cell fitness upon NAM-induced inhibition of Hst3 and Hst4 by influencing Rtt101-Mms1-Mms22-related processes.

Published reports indicate that *hst3*Δ cells display H3 K56ac-dependent defects in the maintenance of a chromosome harboring a reduced number of replication origins (28, 60), which suggests that elevated H3 K56ac might negatively influence the completion of chromosomal DNA replication in situations where the number of active origins is limited. In accord with this, our data indicate that i) cells harboring hypomorphic alleles of the critical origin activation genes *CDC7* and *DBF4* display strong growth defects in the presence of NAM, and ii) firing of early/efficient origins of DNA replication is compromised in *cdc7-4 hst3*Δ *hst4*Δ cells released from G1 toward S at the semi-permissive temperature for *cdc7-4*. Deletion of *RIF1* was found to alleviate these phenotypes, in agreement with the known role of Rif1 in promoting Glc7-dependent dephosphorylation of MCM complexes leading to inhibition of origin activation (33), and with the fact that homozygous deletion of *RIF1* was found to improve cell fitness in response to NAM in our previously published screen (26). Importantly, data presented here also show that lack of Rif1 suppresses several phenotypes of *hst3*Δ *hst4*Δ cells. While these results can be considered surprising in light of the fact that Rif1 has also been reported to promote the stability of stalled RF (41), it is possible that the elevation of origin activity caused by *rif1*Δ overrides the negative impact on replicative stress responses caused by this mutation in *hst3*Δ *hst4*Δ cells.

RF stalling activates Rad53, which then phosphorylates the replication proteins Dbf4 and Sld3 to inhibit the firing of replication origins that have not yet been activated (7). In apparent contrast to our results, a previously published report revealed that *hst3*Δ *hst4*Δ cells present elevated activation of late origins in cells released from G1 toward S phase in medium containing the replication-blocking ribonucleotide reductase inhibitor HU (11). However, such effect was not specific to *hst3*Δ *hst4*Δ mutants; indeed, this was found to be an indirect consequence of elevated spontaneous DNA damage and constitutive Rad53 activity in various replicative stress response mutants, leading to Rad53-dependent elevation of dNTP pools and consequent HU-resistant DNA synthesis (61). In contrast, several observations presented here indicate that constitutive H3 K56ac influences origin firing in a manner that does not depend on ongoing Rad53 activity and/or phosphorylation of Sld3 and Dbf4. First, *cdc7-4 hst3*Δ *hst4*Δ cells do not display noticeable elevation in Rad53 phosphorylation when released from G1 arrest toward S phase at the semi-permissive temperature for *cdc7-4*, even in the presence of HU. Secondly, mutations or treatments that compromise Rad53 activation do not rescue defective S phase progression in *cdc7-4 hst3*Δ *hst4*Δ mutants. Finally, mutations in Sld3 and Dbf4 that abrogate their phosphorylation by Rad53 do not improve DNA replication in *cdc7-4 hst3*Δ *hst4*Δ cells. Taken together, the above data argue that the impact of constitutive H3 K56ac on origin activity in early S phase does not require prior RF stalling and ensuing elevation in Rad53 activity. Consistently, we also showed that the activation of several early/efficient origins is strongly delayed in *cdc7-4 hst3*Δ *hst4*Δ after release from G1 at the semi permissive temperature for *cdc7-4*, which presumably explains why HU exposure does not cause Rad53 activation in these conditions.

Our data and those of others (28, 60) argue that elevated H3 K56ac strongly inhibits DNA replication only under conditions of reduced DDK activity or in situations where the number of active origins is limited. Such conditions are met in *cdc7-4* or *dbf4-1* mutants at the semi-permissive temperature, or in cells harboring an artificial chromosome engineered to have a low number of origin. We emphasize that as mentioned earlier RF stalling leading to Rad53 activation and consequent phosphorylation of Dbf4 also diminishes DDK activity (7). It is therefore possible that genotoxin-induced RF stalling and consequent Rad53 activation might synergize with constitutive H3 K56ac in inhibiting late origin activity in *hst3*Δ *hst4*Δ cells that harbor a wild-type allele of *CDC7*. In turn, this would be expected to compromise the completion of DNA replication, eventually leading to cell death. In agreement with this, we and others previously showed that i) *hst3*Δ *hst4*Δ cells cannot complete DNA replication in a timely manner after transient exposure to genotoxic drugs during S phase (24), ii) *hst3*Δ *hst4*Δ cells present strong and persistent activation of Rad53 upon DNA damage (23, 24, 26), iii) limiting Rad53 activation partially rescues the phenotypes of *hst3*Δ *hst4*Δ mutants (23, 24, 26, 62), and iv) elevating the firing of late origins of replication by overexpression of Cdc45, Sld3 and Sld7 can rescue certain phenotypes caused by elevated H3 K56ac (27, 62). We emphasize that the combined negative effects of Rad53 activation and elevated H3 K56ac on origin activity would be expected to force RF to travel unusually long distances before encountering a converging fork. Consequently, a substantial fraction of persistently stalled RF would not be “rescued” by converging forks in these conditions, leading to under-replicated chromosomal regions, RF collapse, DNA damage, and eventual arrest in G2/M, which are all observed in *hst3*Δ *hst4*Δ cells (12, 19, 23, 24).

The mechanism linking H3 K56ac to origin activity is currently unclear. As mentioned previously, Mec1 activation promotes the degradation of Hst3, which causes inordinate persistence of H3 K56ac in chromatin in cells exposed to genotoxic drugs (57, 58). Such persistence of H3 K56ac in late S might represent a signal that acts redundantly with Rad53 to bolster replicative stress-induced inhibition of late origins of replication. We demonstrated that deletion of *RTT101* or *MMS1*, which encode subunits of a ubiquitin ligase complex previously genetically linked to H3 K56ac (51, 52), rescues the synthetic temperature sensitivity of *hst3*Δ *hst4*Δ *cdc7-4*. Since *rtt101*Δ and *mms1*Δ do not influence H3 K56ac levels (51), any models involving H3 K56ac-dependent modulation of chromatin structure *per se* are therefore unlikely to explain the impact of this modification on origin activity. We note that Rtt101 recruitment to chromatin upon DNA damage was shown to depend at least partly on the H3 K56 acetyltransferase Rtt109 (54). Moreover, Mms1 has been reported to interact directly with the Origin Recognition Complex subunit Orc5 (63), although the biological relevance of this interaction is unclear. The above considerations raise the possibility that DNA damage-induced persistence of H3 K56ac might modulate Rtt101/Mms1 activity *in trans* to downregulate origin activity. Further experiments will be required to test the validity of such models, and to precisely ascertain the mechanistic basis of the impact of H3 K56ac on DNA replication dynamics.

## ACKNOWLEDGEMENTS

H. W. is the recipient of a Chercheur-boursier Sénior scholarship from Fonds de la Recherche du Québec-Santé (award #281795; https://frq.gouv.qc.ca). This work was supported by Natural Sciences and Engineering Research Council of Canada Discovery Grant and Discovery Accelerator Supplement (RGPIN-2019-05082, RGPAS-2019-00009; https://www.nserc-crsng.gc.ca) and by a Fonds de la Recherche du Québec-Nature et Technologies grant (2018-PR-206098; https://frq.gouv.qc.ca) to H.W. C. N. is supported by the Canadian Foundation for Innovation (CFI; https://www.innovation.ca/) and is a Canada Research Chairs (CRC; https://www.chairs-chaires.gc.ca/) Tier 1 Chair. The funders had no role in study design, data collection and analysis, decision to publish, or preparation of the manuscript. We thank Dr David Shore (Université de Genève) for providing yeast strains, Dr Alain Verreault (Université de Montréal) for the generous gift of anti-H3 K56ac and anti-H2A S129-P antibodies, and Dr Elliot A. Drobetsky (Université de Montréal) for critical reading of the manuscript.

## MATERIAL AND METHODS

### Yeast strains and growth conditions

Yeast strains used in this study are listed in Table 1 and were generated and propagated using standard yeast genetics methods. Yeast strains used in Tables S1-S2-S3 were taken from the heterozygote yeast deletion collection (ThermoFisher). For nicotinamide (NAM) treatments, asynchronously growing cells were centrifuged and resuspended at 0.01-0.1 OD/mL in YPD or synthetic (SC) medium containing 20 mM NAM (Sigma-Aldrich). Cells were incubated on a shaker for indicated time. Cells synchronization in G1 was performed by incubating MATa yeasts in medium containing 2 µg/mL alpha-factor for 90 minutes followed by the addition of a second dose of 2 µg/mL of alpha-factor for another 75 minutes. Cells were then washed once in YPD or SC medium and released in S phase in medium supplemented with 5 µg/mL pronase (Protease from *Streptomyces griseus*, Sigma-Aldrich). For ionizing irradiation, exponentially growing cells were exposed to 40 Gy followed by a 60-minute incubation at 30°C prior to sample collection. For methyl methane sulfonate (MMS) treatment, cells were first synchronized in G1 using alpha factor, then incubated in YPD containing 0.01% MMS (Sigma -Aldrich) and 5 µg/mL pronase at a density of 1 OD_630_/mL for 60 minutes. After treatment, cells were washed twice with YPD containing 2.5% sodium thiosulfate (Bioshop), followed by incubation in YPD. Caffeine (Sigma-Aldrich) was used at a concentration of 0.15%.

### Genome-wide fitness screen

The heterozygote diploid yeast fitness screen was realized as described (65–67). Briefly, pools of the yeast hetetozygote diploid deletion mutant collection (BY4743 background) were incubated at 30°C in YPD +/- 41 mM NAM. Cells were collected after 20 generations. PCR reactions were performed on extracted DNA to amplify sequence barcodes, and products were used to probe high-density oligonucleotide Affymetrix TAG4 DNA microarrays. Hybridization, washing, staining, scanning and intensity values calculation were performed as described (65–67). For Z-score calculation, the intensity value of each mutant was divided by the standard deviation. Gene ontology (GO) Term Finder tool was used from the Saccharomyces Genome Database to identify cellular processes affected by NAM treatment (68, 69). Processes identified were considered significant if P-values ≤ 0.01. REViGO was used to summarized significant GO term identified by removing redundant ones (70). Top 1% genes (Z-score > 2.58 or < -2.58) were compared to a previously published screen (performed on homozygote diploid mutants (26)) using Venn diagrams.

### Competitive growth assay

0.0005 OD_630_ of heterozygous deletion and WT diploid yeast cultures were mixed and incubated in YPD +/- 41 mM NAM at 25°C in a 96-well plate. Throughout the incubation, OD_630_ were taken and cells were diluted appropriately to prevent saturation of the culture. After 20 generations, 0.01 OD_630_ of cells was spread on YPD-agar +/- G418 plates. Plates were incubated at 30°C for 48 h and colonies were then counted. The following formula was used to describe growth +/- NAM:

NAM:G418)⁄(NAM:YPD))/((YPD:G418)⁄(YPD:YPD))

NAM:G418 is the number of colonies from cells that were grown in YPD + 41 mM NAM and then plated on YPD + 200 µg/mL G418. NAM:YPD is the number of colonies from cells grown on YPD + 41 mM NAM and then plated on YPD. YPD:G418 is the number of colonies from cells grown on YPD and then plated on YPD + 200 µg/mL G418. YPD:YPD is the number of colonies from cells grown on YPD and then plated on YPD.

### Yeast growth assays

For growth in liquid medium, cells were grown to saturation in YPD in a 96-well plate. Cells were then diluted in fresh medium to 0.0005 OD_630_/ml in 100 µL of YPD containing appropriate concentrations of NAM (Sigma-Aldrich). Cells were then incubated at the indicated temperature for 48-72 h. OD_630_ was then determined using a Biotek EL800 plate reader equipped with Gen5 version 1.05 software. Wells containing YPD were used as blanks. For spot growth assays on solid media, cells were grown in YPD or SC medium in a 96-well plate to equivalent OD_630_. Cells were serially diluted 1:5 and spotted on YPD medium containing nicotinamide (Sigma-Aldrich), methyl methane sulfonate (Sigma-Aldrich), hydroxyurea (BioBasics), or SC medium depleted of uracil (SC-URA) or SC-URA medium containing 50 µg/mL uracil 0.1% of 5-Fluoroorotic acid (Bioshop) (5-FoA). Plates were incubated at the indicated temperature for 48-72 h.

### Cell cycle analysis by flow cytometry

DNA content/cell cycle analysis by flow cytometry was performed as described previously (71). Flow cytometry was performed using a BD Biosciences FACS Calibur instrument equipped with CellQuest software. Data were analyzed using FlowJo 10.8.1 (FlowJo, LLC).

### Immunoblotting

4 OD of cells were pelleted and frozen at -80°C prior whole-cell extraction. Cells were extracted using 0.1M NaOH for 5 minutes at room temperature as described before (72) or using standard tri-chloroacetic acid (TCA) and glass beads method (73). Protein extracts were quantified using bicinchoninic acid (BCA) protein assay kit according to the manufacturer’s protocol (Pierce). SDS-PAGE and transfer were performed using standard methods. Anti-H3 (Abcam; cat: ab1791) and anti-Rad53 (Abcam; cat: ab104232) were purchased from Abcam. Anti-H3 K56ac (AV105) and anti-H2A-S129-P (AV137) antibodies were generously provided by Dr. Alain Verreault (Université de Montréal, Canada). Goat anti-rabbit (BioRad; cat: 1705046), goat anti-mouse (Bio Rad; cat: 1705047) and goat anti-rat (Abcam; cat: ab97057) were used as secondary antibodies. Protein visualization was realized by chemiluminescence using Pierce ECL Western Blotting Substrate. Images were captured using an Azure c600 chemiluminescence Imaging System.

### Fluorescence microscopy

Cells were fixed in 0.1M of potassium phosphate buffer pH 6.4 containing 3.7% formaldehyde (Sigma-Aldrich) and slides were prepared as described (52). Images were taken by fluorescence microscopy using a 60X objective (numerical aperture [NA], 1.42) on DeltaVision instrument (GE Healthcare). Images analysis was performed using SoftWoRx 7 software and FIJI 1.53.

### Pulse-field gel electrophoresis (PFGE)

Yeast chromosome migration by PFGE has been performed as described previously (42). Briefly, 2.5 OD of cells were washed in 50 mM EDTA, then resuspended in 55°C-heated 0.25 mg/mL Zymolyase 100T solution containing 1 % low melting point agarose (LMPA). The mixture is poured into the plug former and placed at 4°C to set. Plugs were then incubated in 0.5 M EDTA, 100mM TRIS-HCl supplemented with 500µL of ß-mercaptoethanol at 37°C overnight followed by two quick washes in 50 mM EDTA. Subsequently, plugs were incubated in 0.325 M EDTA, 1 % N-Lauroylsarcosine, 15 mg proteinase K and 1 mg RNase overnight at 37°C. Plugs were the washed in 1X TE buffer at 4°C for 1h. 1 % agarose gel were prepared in 0.5X TBE buffer. When ready to run, plugs were inserted into the gel. The run is performed at pump setting 70 and at 14°C. Firstly, gel was kept with these parameters for 1h to equilibrate. Migration was performed at 6 V/cm with an angle of 120°. Switch time between pulses were 60 s for the first 15 h, then 90 s for the remaining 9 h. After the migration, gel was stained in H_2_O + 100 µL SYBR Green (Life technologies; cat: S7563) for 1 h. Images were captured using an Azure c600 chemiluminescence Imaging System.

### Alkaline gel electrophoresis and Southern blotting

Samples were denatured by heating at 70°C in loading buffer (30 mM NaOH, 1 mM EDTA, 3% Ficoll 400, 0.01% bromocresol green). Denatured DNA was run in a 1% agarose gel in alkaline electrophoresis buffer (30 mM NaOH, 2 mM EDTA) at 3 V/cm. Southern blotting was performed using a digoxygenin (DIG)-labeled probe as described (74). The ARS305 probe was generated by PCR using primers ARS305_probe_F and ARS305_probe_R (Table 1) and the PCR DIG Labeling Mix (Roche). Membranes were imaged using an Azure c600 chemiluminescence Imaging System.

### Mesurement of BrdU incorporation by DNA immunopurification and qPCR

Measurement of BrdU signal was performed as described in (75). Briefly, 400 µg/ml BrdU (BioShop; cat: BRU222.5) was added to cells release toward S phase at 37°C for 1h. 10 OD of cells were harvested per condition. Cells were immediately fixed in 70% ethanol. DNA was extracted as described (76), then sonicated at 25 % for four cycles of 15 s. DNA samples were then purified using EZ-10 Spin Column PCR Products Purification Kit (Bio Basics) according to the manufacturer instructions. DNA was quantified using a fluorometer (Turner Biosystems) according to the manufacturer’s protocol. 500ng of genomic DNA was used per condition for immunoprecipitation. DNA samples were mixed with 0.1 µg/µL of blocking DNA and denaturated at 95°C for 10 minutes followed by snap-cooling on ice for 5 minutes. Then, DNA samples were incubated with anti-BrdU antibody (Invitrogen; cat: ZBU30) antibody in 1X PBS + 0.0625 % Triton X-100 at 4°C for 4 h. 15 µL of washed Protein G MagBeads (GenScript; cat: L00274) were added to the DNA samples and incubated overnight at 4°C. Immunoprecipitated DNA samples were washed 3 times with 500 µL 1X PBS + 0.0625 % Triton X-100 and two times with 1X TE pH 7.6. Then, samples were eluted with TE pH 7.6 1 % sodium dodecyl sulfate (SDS) at 65°C for 15 minutes. The eluate is then purified using EZ-10 Spin Column PCR Products Purification Kit (Bio Basics) according to manufacturer’s instructions. 3 µL of immunoprecipitated sample or 0.5 ng of input sample was used per qPCR reaction (qPCR Master Mix, APExBio; cat: K1070). PCR was performed using an Applied Biosystems 7500 instrument (software version 2.3). PCR primers are listed in Table 1. BrdU incorporation quantification were performed using the standard percent of the input method. To normalize for eventual differences in BrdU incorporation capacity between strains, IP/input value was divided by the IP/input value obtained using the same strain which was synchronized in G1 and then released in S phase for 1 h in presence of 200 mM HU at 25°C.

### Measurement of DNA content by quantitative PCR

Genomic DNA from 1 OD_630_ of cells was extracted and purified as described (76). 3 ng of DNA was used per qPCR reaction (qPCR Master Mix, NEB). PCR was performed using an Applied Biosystems 7500 instrument (software version 2.3). PCR primers are listed in Table 1. Briefly, qPCR signal for a given origin was first normalized to the signal obtained from the NegV locus (ChrV: 532538-532516) (62). This region is located ≈12 Kb from ARS521, an origin which has not been detected to be active in several studies according to OriDB (http://cerevisiae.oridb.org/) and ≈18 kb from ARS522, a subtelomeric origin of replication activated in late S. As such, the NegV locus is expected to be replicated in late S, and therefore to generally remain unreplicated in a majority of *cdc7-4* and *cdc7-4 hst3*Δ *hst4*Δ cells 30 minutes post-release from G1 arrest toward S phase. The NegV-normalized S phase signal was divided by the NegV-normalized signal obtained from alpha factor arrested (G1) cells. Complete replication of an origin is therefore expected to result in a ratio of S phase over G1 signal of 2.

### Statistical analysis

Data are represented as mean ± standard error of the mean (SEM) unless otherwise specified. All analyses were performed using GraphPad Prism 8. Statistical tests used are described in figure legends.

