## Supplementary material for "Persistent acetylation of histone H3 lysine 56 compromises the activity of DNA replication origins": Table S2 and S3

### Table S2. GO term analysis of gene with positive Z-score in NAM fitness assay

| Gene Ontology term | Cluster frequency | Genome frequency | Corrected P-value | FDR | False Positives | Genes annotated to the term |
|---|---|---|---|---|---|---|
| macromolecule metabolic process | 98 of 131 genes, 74.8% | 3217 of 7166 genes, 44.9% | 1.76e-09 | 0.00% | 0.00 | PSY2, PPH3, RFA2, BSD2, MAK10, GFD1, YKU80, MRC1, ALG2, PRP9, TAF10, SMT3, RRP4, KRE29, DBF2, RPN13, UTP15, NPP1, RPL32, PKC1, YKU70, MTR3, MNL1, TIF11, SMX2, GSP1, HYP2, TFC1, CUS1, PSF2, CFT1, SLD2, TAF5, TAF6, PIN4, RRP45, WRS1, SPN1, MPS3, SLD3, PDC2, UBP3, ERD1, TOF1, SGT2, SRB7, PUT3, UFD4, PRI2, YHC1, ACA1, POL32, TIF2, NOP58, TFB1, ASC1, MED11, MCM4, LSM2, DBF4, MCD4, GLC7, FPR3, NUP145, TAF7, RRP42, RFA1, SSL2, SRS2, BRE5, NMD4, SEN1, DBP2, KAP95, KKQ8, STE11, DUN1, SLX4, CDC12, TAF8, NIP1, MAK31, UBC9, NET1, DPB3, KEX2, NUP82, SSN3, TIP41, LAS1, DPB4, ORC3, SPT15, UBP8, BDP1, SUI1, SAS10, TAF12 |
| DNA-dependent DNA replication | 18 of 131 genes, 13.7% | 133 of 7166 genes, 1.9% | 1.78e-08 | 0.00% | 0.00 | DPB3, SLD2, MCM4, DBF4, MRC1, POL32, PSF2, PRI2, RFA2, DUN1, SLX4, TOF1, SEN1, RFA1, SLD3, GLC7, ORC3, DPB4 |
| cellular response to DNA damage stimulus | 27 of 131 genes, 20.6% | 357 of 7166 genes, 5.0% | 1.37e-07 | 0.00% | 0.00 | PSY2, PPH3, RFA2, DUN1, SLX4, IRC21, PRI2, YKU80, PSF2, POL32, MRC1, TFB1, SLD2, DPB3, MCM4, KRE29, GLC7, NUP145, DPB4, PIN4, RFA1, SLD3, SSL2, SRS2, SEN1, TOF1, YKU70 |
| DNA integrity checkpoint | 10 of 131 genes, 7.6% | 55 of 7166 genes, 0.8% | 3.53e-05 | 0.00% | 0.00 | PIN4, PPH3, PSY2, DUN1, GLC7, SLX4, SLD2, TOF1, DBF4, MRC1 |
| DNA geometric change | 11 of 131 genes, 8.4% | 71 of 7166 genes, 1.0% | 4.14e-05 | 0.00% | 0.00 | MCM4, YKU70, SLD2, SRS2, SEN1, YKU80, PSF2, RFA1, SSL2, SLD3, RFA2 |
| biological regulation | 66 of 131 genes, 50.4% | 2096 of 7166 genes, 29.2% | 0.00019 | 0.00% | 0.00 | DFR1, GSP1, HYP2, SLD2, TAF6, PIN4, RRP45, SPN1, MPS3, PDC2, UBP3, ERD1, TOF1, SRB7, SGT2, PUT3, PSY2, PPH3, RFA2, BSD2, YKU80, MRC1, RRP4, DBF2, ZRC1, RPN13, UTP15, PKC1, MTR3, YKU70, KAP95, KKQ8, STE11, DUN1, SLX4, CDC12, NET1, DPB3, SSN3, TIP41, DPB4, ORC3, SPT15, BDP1, SUI1, UFD4, ECM25, YHC1, ACA1, TFB1, MED11, ASC1, DBF4, FPR3, GLC7, NUP145, TAF7, RRP42, RFA1, SSL2, SRS2, BRE5, SEN1, NMD4, FTH1, DBP2 |
| cellular component organization or biogenesis | 73 of 131 genes, 55.7% | 2453 of 7166 genes, 34.2% | 0.00024 | 0.00% | 0.00 | SPT15, ATP11, LAS1, DPB4, ORC3, TIM10, SAS10, TAF12, SUI1, BDP1, UBP8, CDC12, SLX4, KAP95, NUP82, DPB3, UBC9, NET1, NIP1, TAF8, RFA1, SSL2, RRP42, GLC7, FPR3, TAF7, NUP145, MCD4, DBP2, SRS2, SEN1, ECM25, YHC1, UFD4, VTC3, MED11, MCM4, LSM2, DBF4, CDC3, NOP58, SPN1, MPS3, SLD3, RRP45, PUT3, TOF1, HYP2, TFC1, GSP1, TIF11, SMX2, TAF6, TAF5, SLD2, PSF2, CUS1, DBF2, TAF10, RRP4, TIM12, YJL160C, PKC1, YKU70, MTR3, UTP15, RFA2, ECM30, PPH3, PSY2, PRP9, YKU80, MRC1 |
| cellular process | 117 of 131 genes, 89.3% | 5175 of 7166 genes, 72.2% | 0.00113 | 0.00% | 0.00 | TIF11, MNL1, DFR1, SMX2, GSP1, TFC1, HYP2, PSF2, CUS1, CFT1, SLD2, TAF5, TAF6, DPP1, NKP1, PIN4, CAR2, WRS1, RRP45, SPN1, MPS3, SLD3, UBP3, PDC2, ERD1, TOF1, SGT2, SRB7, PUT3, PPH3, PSY2, RFA2, ECM30, BSD2, IRC21, MAK10, YKU80, MRC1, ALG2, PRP9, SMT3, TAF10, RRP4, KRE29, DBF2, ZRC1, HOM6, RPN13, UTP15, RPL32, NPP1, CCT3, YJL160C, TIM12, PKC1, YKU70, MTR3, CCT6, KKQ8, KAP95, STE11, DUN1, SLX4, SAM2, CDC12, TAF8, NIP1, MAK31, UBC9, NET1, DPB3, SSN3, TIM10, TIP41, LAS1, ORC3, DPB4, SPT15, ATP11, DOG1, BDP1, UBP8, SUI1, SAS10, TAF12, UFD4, VTC3, ECM25, PRI2, YHC1, ACA1, POL32, MIS1, NOP58, TIF2, CDC3, TFB1, ASC1, MED11, MCM4, LSM2, DBF4, MCD4, GLC7, FPR3, TAF7, NUP145, RRP42, RFA1, TCP1, SSL2, BRE5, SRS2, SEN1, NMD4, CCT2, DBP2 |

### Table S3. GO term analysis of genes with negative Z-score in NAM fitness assay

| Gene Ontology term | Cluster frequency | Genome frequency | Corrected P-value | FDR | False Positives | Genes annotated to the term |
|---|---|---|---|---|---|---|
| ubiquitin-dependent protein catabolic process | 18 of 58 genes, 31.0% | 226 of 7166 genes, 3.2% | 2.46e-11 | 0.00% | 0.00 | RPT2, TRE2, SSE1, SCL1, PRE7, PRE8, RPT3, RPN5, RPN11, MET30, PRE3, RPT1, RPN12, PRE1, PUP2, PRE6, RPT5, RPN6 |
| proteasome assembly | 7 of 58 genes, 12.1% | 35 of 7166 genes, 0.5% | 3.04e-06 | 0.00% | 0.00 | RPT1, RPN6, RPN11, RPT5, PNO1, RPT2, RPT3 |
| proteasome regulatory particle assembly | 4 of 58 genes, 6.9% | 11 of 7166 genes, 0.2% | 0.00042 | 0.00% | 0.00 | RPT1, RPT2, RPT3, RPT5 |
| catabolic process | 21 of 58 genes, 36.2% | 936 of 7166 genes, 13.1% | 0.00211 | 0.00% | 0.00 | RPT2, TRE2, SSE1, SCL1, PRE7, PRE8, RPT3, RPN5, LSM7, RPN11, MET30, RPL13B, CDC39, RPN12, RPT1, PRE3, PRE1, PRE6, PUP2, RPT5, RPN6 |
